# Universal dynamics of biological pattern formation in spatio-temporal morphogen variations

**DOI:** 10.1101/2022.03.18.484904

**Authors:** Mohit P. Dalwadi, Philip Pearce

**Affiliations:** Department of Mathematics, University College London; Institute for the Physics of Living Systems, University College London

## Abstract

In biological systems, chemical signals termed morphogens self-organise into patterns that are vital for many physiological processes. As observed by Turing in 1952, these patterns are in a state of continual development, and are usually transitioning from one pattern into another. How do cells robustly decode these spatio-temporal patterns into signals in the presence of confounding effects caused by unpredictable or heterogeneous environments? Here, we answer this question by developing a general theory of pattern formation in spatio-temporal variations of ‘pre-pattern’ morphogens, which determine gene-regulatory network parameters. Through mathematical analysis, we identify universal dynamical regimes that apply to wide classes of biological systems. We apply our theory to two paradigmatic pattern-forming systems, and predict that they are robust with respect to non-physiological morphogen variations. More broadly, our theoretical framework provides a general approach to classify the emergent dynamics of pattern-forming systems based on how the bifurcations in their governing equations are traversed.

## 1 Introduction

In biological pattern formation, cells interpret morphogen signals to make developmental decisions based on their location in a tissue, organ, embryo or population. Such systems have been found to be remarkably robust to a wide range of sources of variation in morphogen signals and system components [1–3]. For example, in recent work, general principles of biological pattern-forming systems have been identified that promote robustness with respect to variations in morphogen and protein production rates [2, 4], in tissue or organism size [5, 6], and in gene-regulatory network architecture [7]. However, these studies have generally focused on robustness with respect to variations from cell to cell or from tissue to tissue [2] – much less is understood about how such systems respond to spatio-temporal variations in morphogen concentrations within individual cells, populations or tissues, particularly over timescales faster than or similar to growth.

Recent experimental and theoretical work has demonstrated how specific gene-regulatory network architectures convert spatio-temporal morphogen signals into a required static or dynamic response [8–13]. These morphogen signals, which have been called ‘pre-pattern’ morphogens, can arise as a necessary part of the developmental process [8, 14, 15]. However, unpredictable pre-pattern morphogen fluctuations in a system may be caused by intrinsic noise [16], growth [17], cell motility or rearrangement [18–21], biochemical reactions [22, 23], or external flows [12, 24–26]. These studies raise the question of how to quantify the robustness of a system’s gene-regulatory network output, i.e. emergent spatio-temporal patterning, with respect to variations in its pre-pattern morphogen input over a certain timescale. A typical approach to mathematical modelling of biological pattern formation is to use reaction-diffusion equations that capture the essential dynamics of relevant morphogens [8]. In reaction-diffusion equations, gene-regulatory networks are represented using the reaction terms, and pre-pattern morphogens can be taken into account through the associated reaction parameters [15]. Such equations display self-organisation in space and time via bifurcations [27, 28], which often cause switch-like transitions in the solution that may correspond to biological decisions [12] or cell-fate choices [29]. Mathematically, dynamics are typically observed to slow down near bifurcations [30, 31], and these ‘delayed bifurcation’ effects have previously been considered in reaction-diffusion contexts [32–34], but the implications to biological pattern formation and decision-making are not understood.

Here, we connect bifurcations in reaction-diffusion systems to robustness in biological pattern formation. Although our approach is general, we consider two specific pattern-forming systems with spatio-temporal variations in pre-pattern morphogens. Mathematically, the morphogen variations correspond to variations in the parameters of the governing reaction-diffusion equations. We use mathematical analysis to classify and quantify the dynamic response to such variations in terms of universal solution regimes that we identify. The regimes emerge as a consequence of bifurcations in the governing equations – in corresponding biological systems, the type of bifurcation is determined by the system’s gene-regulatory network. Our analysis predicts an emergent timescale *associated with bifurcations*; in the relevant solution regime, this bifurcation timescale allows the system to filter out oscillatory variations in pre-pattern morphogens that occur faster than a critical timescale. Surprisingly, in the two biological examples that we study, we find that the critical timescale is a few hours shorter than physiological timescales. Overall, our analysis and simulations suggest that gene-regulatory networks in some biological systems may have been tuned for robustness: they are able ignore to variations or oscillations in morphogen concentrations that happen much faster than growth, while responding appropriately to physiological morphogen variations.

The paper contains three main parts. In the first part, we derive (Section 2) and analyse (Section 3) a generic canonical equation, or normal form, that captures pattern-formation dynamics near supercritical pitchfork and transcritical bifurcations. In the second part, we model and analyse two specific biological systems so that they can be written in terms of the canonical results (Section 4 and Appendix C). In the final part, we summarise our results and discuss their implications for biological pattern formation (Section 5 and Section 6). Therefore, the reader most interested in the biological applications of this study can go directly to Section 5.

## 2 Derivation of canonical equations

As a reaction-diffusion system undergoes a bifurcation, its dynamics can typically be approximated by a (lowdimensional) canonical weakly nonlinear equation whose form depends on the type of bifurcation [30]. To perform analysis relevant to several classes of biological pattern formation, we here focus on such a canonical equation that incorporates pitchfork and transcritical bifurcations. We then specialise to quantify dynamics as system parameters are varied; the dynamics depend on the type of bifurcation, and in which direction the system passes through the bifurcation (either ‘on-to-off’ or ‘off-to-on’). As will be made clear in Section 5, each type of bifurcation that we study can be associated with a specific biological pattern-forming system, and parameter variations can be linked to a specific pre-pattern morphogen.

Under a weakly nonlinear approximation, the effects of spatio-temporal variations in system parameters are captured by a specific variation in the parameters of the related weakly nonlinear equation near the bifurcation. Moreover, we will show that variations that may appear quasi-steady away from the bifurcation can depend on the dynamics of the crossing near the bifurcation. This dynamic effect can result in an effective robustness in the system over non-standard timescales.

To understand this effect, we first demonstrate how the appropriate inner canonical equation, or normal form, can emerge in general through a systematic asymptotic analysis in the ‘inner region’ near the bifurcation. To this end, we first consider the effect of spatio-temporal variations in the parameters of the dimensionless toy reaction-diffusion problem

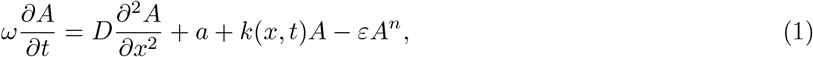

where we have non-dimensionalised time and space over their typical scales in the spatio-temporal variation. In Eq. (1), *A*(*x, t*) can be thought of as a measure of the morphogen concentration (in terms of the deviation from the non-patterned state for the mode excited over the bifurcation^1^). The parameter *D >* 0 represents a strength of diffusion. The parameter *a* ≥ 0 represents a measure of base production in terms of the excited mode; *a* typically vanishes for noiseless Turing systems (*n* = 3). The imposed function *k*(*x, t*) represents the net strength of self-activation in comparison to decay. When *k >* 0, the net effect is self-activation, while when *k <* 0 the net effect is decay. The parameter *ω >* 0 represents a measure of the frequency of *k* variation and *ε >* 0 represents the strength of nonlinear saturation effects. The exponent *n* = 2 or 3, corresponds to the canonical weakly nonlinear form of an imperfect transcritical or a (supercritical) pitchfork bifurcation, respectively. Each of these characterises a minimal gene-regulatory motif (Fig. 1a). The case with *n* = 2 (*n* = 3) can be thought of as a modified version of the Fisher-KPP (Ginzburg-Landau) equation. Generally, to obtain the significant ‘on-off’ type effects seen in biological signalling, the saturation effect must be weak i.e. *ε* ≪ 1. For ease of exposition, it is also convenient to impose *ω* ≪ 1 and *D* ≪ 1, so that spatio-temporal variations in *k* appear to be locally quasi-steady in (1). We will show that, contrary to its appearance, the system (1) is not generally locally quasi-steady when crossing a bifurcation, and that this has significant implications for robustness over non-standard timescales. At this point, we make no further assumptions on the relative sizes of the small parameters *ε, ω*, and *D*; we seek to understand all possible significant changes in system behaviour as the relative sizes of these parameters vary within the constraint of them being small.

**Figure 1:**
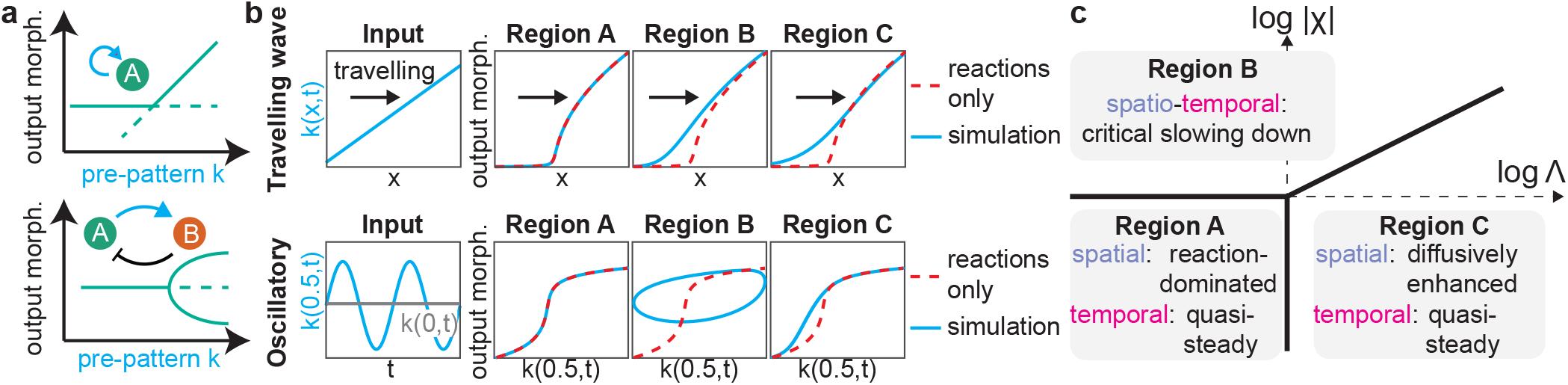
General asymptotic framework for classifying dynamics of pattern-forming systems. a) Bifurcation diagrams and related minimal gene-regulatory network motifs for two classes of pattern-forming system. Top: autoinduction loops are associated with transcritical bifurcations. Bottom: activator-inhibitor systems are associated with pitchfork bifurcations. b) Top: Examples of predicted dynamics in three different regions of parameter space (see panel c) for a travelling gradient, the non-degenerate linearized form of more general spatio-temporal variations. Results were generated by solving Eq. (1) numerically, with *n* = 3 and *k*(*x, t*) specified via *k*(*x, t*) = *x* − *t*. Results are similar for *n* = 2. The dashed line shows reaction-dominated results obtained by solving the steady form of Eq. (1) for each *x* without diffusion. Bottom: Examples of predicted dynamics for a spatio-temporal oscillation in *k*(*x, t*) of the form *k*(*x, t*) = − 0.475 + *x* (1 + 0.25 sin(2*πt*)), shown at a fixed *x* (left). Results were generated as in the top row, and plotted at *x* = 0.5 in a domain of size *x* = 1 (right; y axis log scale). In Region B, the system becomes locked in the patterned state despite large oscillations in *k*(*x, t*). c) General parameter space that classifies the system dynamics in each region for a general spatiotemporal variation. The boundary between Regions B and C occurs when Λ^2*/*3^*/*|*χ*| = *O* (1) and Λ, |*χ*| ≫ 1.

Given the above, in the apparent locally quasi-steady system, *k <* 0 represents the unpatterned region (where *A* ∼ *a/*(−*k*) = *O* (1)) and *k >* 0 the patterned region (where *A*^*n*−1^ ∼ *k/ε* ≫ 1), with *k* = 0 defining the position of the bifurcation in homogeneous conditions. Immediately we can see that these locally quasi-steady solutions will not hold near *k* = 0, and this leads to the natural definition of the moving position *x* = *s*(*t*) of the bifurcation in homogeneous conditions, defined through *k*(*s*(*t*), *t*) = 0. By examining the behaviour of the system near the bifurcation at *x* = *s*(*t*), we will see how and when robustness can be generated in the system.

### 2.1 Canonical inner problem

To derive the canonical inner problem, we must transform into the frame around the moving point *x* = *s*(*t*), at which *k* = 0. This focuses our attention onto the local region near the moving bifurcation. For a travellinggradient-like motion (i.e. a non-degenerate bifurcation moving monotonically), we have *η*(*t*) := *k*_*x*_(*s*(*t*), *t*) ≠ 0 and 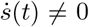, and this is the scenario we focus on below. Later, in Section 3.2, we consider the scenario with oscillating parameters when the bifurcation turns around. Finally, while not intrinsically necessary, we note it is convenient to treat the cases of imperfect and perfect bifurcations separately.

#### 2.1.1 Imperfect bifurcations

For imperfect bifurcations, we introduce the scaled variable *Z*(*x, t*) = *O* (1), defined through

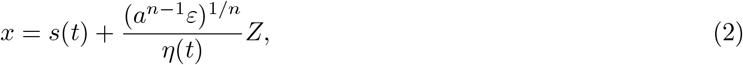

emphasising that *ε* ≪ 1, so we are zooming into a region near *x* = *s*(*t*). From the definition (2), derivatives are transformed as follows

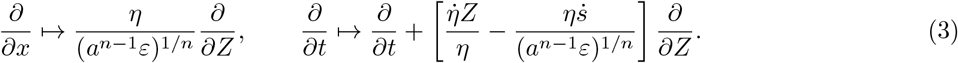

Since *s*(*t*) is defined such that *k*(*s*(*t*), *t*) = 0, by substituting (2) into *k*(*x, t*) and Taylor expanding, we find that

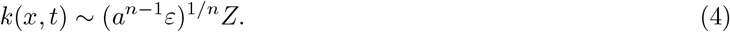

Substituting (3)–(4) into (1), and using the dependent variable transform *Y* (*Z, t*) = (*ε/a*)^1*/n*^*A*(*x, t*), we obtain the transformed system

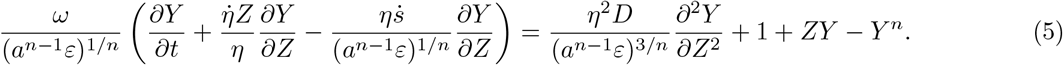

Given that *ε* ≪ 1 and *η*, 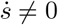 for the non-degenerate moving bifurcation case we consider, the first two terms in the brackets on the left-hand side are asymptotically sub-dominant to the third. Since we have made no further assumptions on the relative sizes of *ε, ω*, and *D* beyond them being intrinsically small, we may consistently retain all remaining terms in order to capture as much information as possible, which represents a distinguished asymptotic limit of the system. This procedure yields the leading-order system

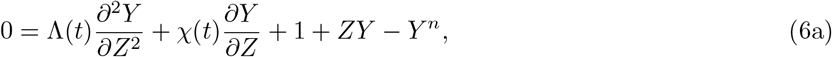

where

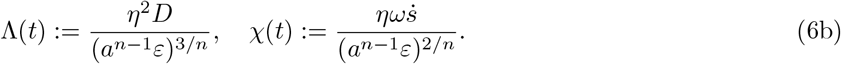

Broadly, Λ ∈ (0, ∞) quantifies the importance of diffusion and *χ* ∈ (−∞, ∞) quantifies the importance of spatio-temporal changes in the parameters, both in comparison to base production. The sign of *χ* relates to whether the bifurcation is moving into the patterned or unpatterned region; the difference between these cases can be mathematically and biologically significant, as we will discuss later.

Finally, we note that the appropriate far-field conditions can be obtained by matching into the outer (positive) quasi-steady solutions. In terms of the canonical equation (6), the consistent matching conditions are

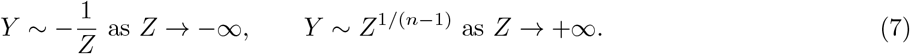

#### 2.1.2 Perfect bifurcations

For a perfect bifurcation, *a* = 0 in (1). Although this case can be treated through appropriate rescalings and limits of the imperfect case, it is more straightforward to simply rescale differently from the outset. Hence, for a perfect bifurcation we introduce the scaled variable *z*(*x, t*) = *O* (1), via

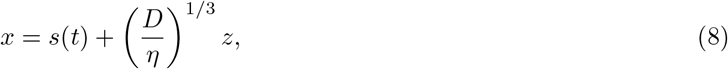

emphasizing that *D* ≪ 1, so we are zooming into a region near *x* = *s*(*t*). Under equivalent reasoning as for the imperfect case, noting that *k*(*x, t*) ∼ *D*^1*/*3^*η*^2*/*3^*z*, and using the dependent variable transform *y* = (*D*^1*/*3^*η*^2*/*3^*/ε*)^1*/*(*n*−1)^*A*, we obtain the transformed system

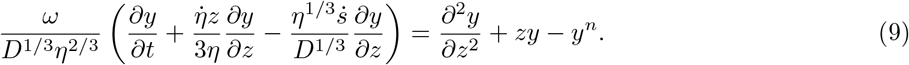

Similarly to before, but now exploiting *D* ≪ 1, for non-degenerate cases the first two terms in the brackets on the left-hand side are asymptotically sub-dominant to the third. Consistently retaining all remaining terms represents a distinguished asymptotic limit of the leading-order system, yielding

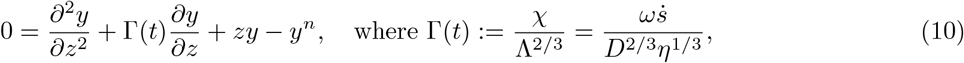

noting that Λ and *χ* are defined in (6b), and Γ ∈ (−∞, ∞) quantifies the importance of spatio-temporal variations in comparison to diffusion. The sign of Γ relates to whether the bifurcation is moving into the patterned or unpatterned region.

The appropriate far-field conditions can be obtained by matching into the outer quasi-steady solutions. Although there is some subtlety involved in matching into the unpatterned region in general for perfect bifurcations, in terms of what is required for our purposes we may close the canonical equation (6) by writing our matching conditions as

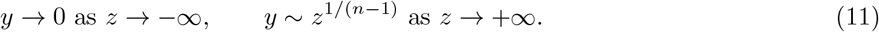

## 3 Asymptotic analysis of canonical system

To fully characterize the possible behaviours of the transition across the bifurcation, we seek to understand the possible behaviours of *C* in terms of Λ and *χ*. One way to do this is to examine the distinguished asymptotic limits of the ODE (6) with boundary conditions (7). This approach has the benefit that we will be able to derive analytic leading-order solutions, which will allow us to quantify the effect of crossing the bifurcation.

We now briefly summarise the asymptotic structure of the solution space we will explore below, before presenting the analysis. As outlined in Fig. 1c, in terms of asymptotic values of Λ and *χ* there are three sublimits in the system labelled Regions A-C, separated by distinguished limits. Each sublimit corresponds to the overall domination of one of the three physical drivers within the transition region: positive feedback, diffusion, and spatio-temporal variations in the parameters. Given that we are interested in understanding how the solution varies away from the quasi-steady solution dominated by positive feedback, it is helpful to define the point *Z* = *Z*\* at which *C*(*Z*\*) = 1. We refer to this as the dynamical position of the bifurcation. We will show below that *Z*\* = 0 for the straightforward quasi-steady solution, but that *Z*\* ≠ 0 when diffusive and spatio-temporal variations are important in the system.

For pitchfork bifurcations (*n* = 3) arising from activator-inhibitor systems in e.g. Turing patterns, a key question is whether the patterned regions remain robust under spatio-temporal variations involving incursions from the unpatterned state (‘on-to-off’). This corresponds to the bifurcation moving from the unpatterned to the patterned state (*χ >* 0). For transcritical bifurcations (*n* = 2) arising from self-activator systems, a key question is whether unpatterned regions remain robust under spatio-temporal variations involving incursions from the patterned state (‘off-to-on’). This corresponds to the bifurcation moving from the patterned to the unpatterned state (*χ <* 0). While the specific details of the analysis will depend on whether the bifurcation is moving towards the patterned or the unpatterned region, the appropriate scalings for the distinguished limits and sublimits outlined in Fig. 1c hold in either case. In an effort to balance breadth and brevity of analysis, we therefore limit ourselves here to considering the two physically relevant cases mentioned above. We present the pitchfork case (*n* = 3) with *χ >* 0 in the main text below, and the transcritical case (*n* = 2) with *χ <* 0 in Appendix B.

In the following, we first analyse travelling-wave-like variation of the system parameters (i.e. monotonic, unidirectional motion of the imperfect bifurcation), and then analyse the scenario with spatio-temporally oscillating parameters (i.e. where the bifurcation turns around). Together, these encompass a significant class of behaviours the moving bifurcation can exhibit.

### 3.1 Travelling-wave-like parameter variations for on-to-off pitchfork bifurcations

In this subsection we consider pitchfork bifurcations (*n* = 3), where the bifurcation moves unidirectionally from the unpatterned to patterned state (*χ >* 0). This corresponds to ‘on-to-off’ patterning. The imperfect pitchfork analysis involves the governing equation (6), taking *n* = 3, with far-field conditions (7). Although the perfect pitchfork analysis involves the governing equation (10) with *n* = 3 and far-field conditions (11), much of the analysis is the same as for the imperfect case. We therefore present the imperfect case in its entirety and flag any differences with the perfect case when they arise.

We re-emphasize that in this Section we are specifically interested in asymptotic solutions for *χ >* 0, and the analytic results we obtain for *C* and *Z*\* are specific to this case. However, the asymptotic scalings we derive here for each sublimit, and summarise later in Section 3.1.4, will be the same for *χ <* 0. We proceed by analysing each region in increasing order of complexity.

#### 3.1.1 Positive feedback dominates: Region A

The simplest region is Region A, where Λ, *χ* ≪ 1. Hence, positive feedback dominates here, and the solution remains quasi-steady through the bifurcation. In this case, the ODE (6) becomes quasi-steady, and at leading-order *Y* is defined implicitly through the cubic equation

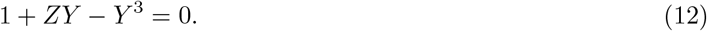

We note that, if required, an explicit solution to (12) can be obtained straightforwardly by invoking the cubic formula. In this quasi-steady limit, *Y* (0) = 1, and therefore *Z*\* = 0. Hence, in Region A the bifurcation takes its equilibrium position as we would expect from a quasi-steady analysis, and the patterning is spatially local and quasi-steady.

Since positive feedback dominates and balances base production here, this region is not relevant in the case of a perfect bifurcation.

The solutions in the remainder of parameter space will be significantly different to Region A. In particular, there will be sharp transitions between the unpatterned and patterned parts of the transition region, and in general these transitions will not occur around *Z* = 0.

#### 3.1.2 Spatio-temporal variations dominate: Region B

Region B occurs when *χ* ≫ 1 and Λ*/χ*^3*/*2^ ≪ 1. In this region, spatio-temporal changes in the parameters dominate and delaying effects become important, with implications for pattern robustness. Importantly, in this region there are sharp transitions between the unpatterned and patterned parts of the transition region, separated by *Z* = *Z*_*c*_ (which we shall see is different but related to *Z*\*).

To summarise the asymptotic structure of the solution before going into the specific mathematical analysis - in Region B, the interesting behaviour occurs over the lengthscale 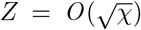, so we scale into a new independent variable 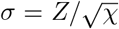. The asymptotic solution is then split into two asymptotic regions separated by the point 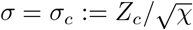. The determination of *σ*_*c*_ and *Z*\* are key goals of our analysis. The unpatterned region corresponds to *σ < σ*_*c*_ and the patterned region corresponds to *σ > σ*_*c*_. The solution is smaller in the unpatterned region, with the scaling 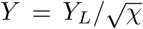, and larger in the patterned region, with the scaling *Y* = *χ*^1*/*4^*Y*_*R*_.

Here, information travels in the same direction as the pattern transition i.e. from the patterned state to the unpatterned state (on-to-off). As such, our analysis starts in the patterned state, where *σ > σ*_*c*_. Using the scalings noted above, the leading-order scaled version of the ODE system (6), (7) in the patterned state is

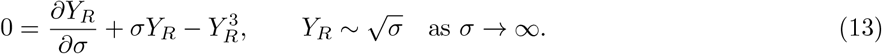

We can solve (13) explicitly by multiplying through by *Y*_*R*_, and introducing the new dependent variable *W* (*σ*) such that 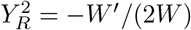. The ODE (13) then transforms to

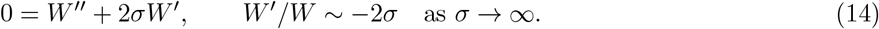

The two linearly independent solutions to the ODE in (14) are erfc(*σ*) and 1, where erfc is the complementary error function. Applying the far-field condition in (14) removes the possibility of any contribution from the constant solution, leading to the solution *W* = *α* erfc(*σ*) for constant *α*, and hence

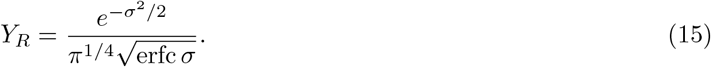

For later matching purposes, we note the following far-field result from (15):

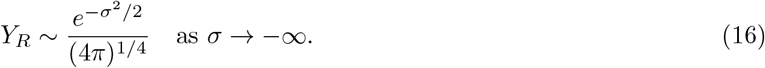

Given the relative scalings of *Y* in the patterned and unpatterned regions, we can determine *σ*_*c*_ by calculating when *Y*_*R*_ = *O* (*χ*^−3*/*4^). From the far-field result (16), this occurs when 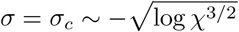.

For completeness, we now also provide the solution in the unpatterned state, where *σ < σ*_*c*_. Using the scalings noted above for the unpatterned state, the leading-order scaled version of the ODE system (6), (7) in the unpatterned state is

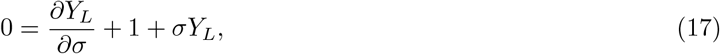

with an additional condition that will arise from matching with the solution (15) in the patterned state. Using the integrating factor method, the ODE (17) is solved by

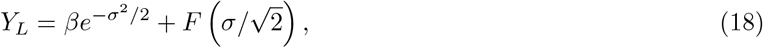

where *F* (*x*) is Dawson’s integral [35, Eq. 7.2.5], defined as

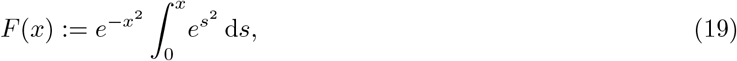

and *β* is a constant to be determined through matching with (15). Since Dawson’s integral (19) is bounded along the entire real line, the matching procedure does not involve this term and is therefore fairly straightforward. Using the far-field limit of the patterned solution (16), and recalling that the intrinsic solution scalings are 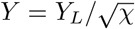 and *Y* = *χ*^1*/*4^*Y*_*R*_, we find that

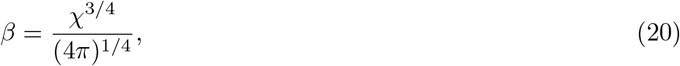

completing the solution (18).

Finally, we calculate the dynamic position of the bifurcation 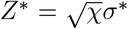, defined through *Y* (*Z*\*) = 1 and hence through *Y*_*R*_(*σ*\*) = *χ*^−1*/*4^. From (16), this occurs for 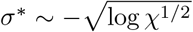. Scaling back into *Z*\*, we obtain the following asymptotic result for the dynamic position of the bifurcation:

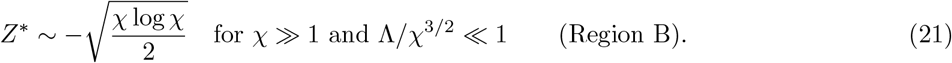

For a perfect pitchfork, as defined in (10), (11), the Region B sublimit is equivalent to Γ ≫ 1. In this case, the scalings *z* ∼ Γ^1*/*2^ and *y* ∼ Γ^1*/*4^ lead to the ODE (13) at leading order, so the appropriate solution is the equivalent of (15):

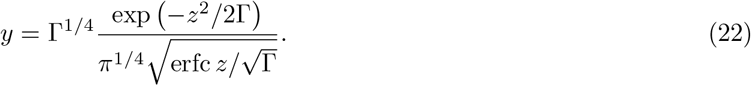

However, for a perfect bifurcation there is no base production to mediate the exponential decay in the far-field as *z* → −∞. Hence, dependent on the actual far-field matching conditions (which will depend on the specific system and its history), it is possible for diffusive effects to reassert themselves in the far-field. That is, there is an additional feasible asymptotic region in this sublimit when −*z* = *O* (Γ^2^). In this region, *y* is exponentially small and the appropriate version of (10) is its linearization, so that saturation effects can be ignored. A WKBJ analysis of the linearization of (10) (matching appropriately with (22)) yields the solution

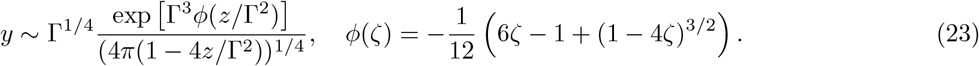

Hence, the position of the dynamic bifurcation (and the extent of the transition region) for the perfect case will depend on how small one requires *y* to be when defining the position of the dynamic bifurcation. Our results above allow for straightforward calculation of the extent once this critical size is defined.

#### 3.1.3 Diffusion dominates: Region C

Region C occurs when Λ ≫ 1 and Λ*/χ*^3*/*2^ ≫ 1. In this region, diffusive effects dominate. In a similar manner as for Region B, there are sharp transitions between the unpatterned and patterned parts of the transition region here, separated by *Z* = *Z*_*c*_ (which again is different but related to *Z*\*).

To summarise the asymptotic structure of the solution - in Region C the interesting behaviour occurs over the lengthscale *Z* = *O* (Λ^1*/*3^), so we scale into a new independent variable *ζ* = *Z/*Λ^1*/*3^ = *O* (1). The solution is again split into two asymptotic regions, matched using an intermediate transition region. This time, the two main asymptotic regions are separated by the point *ζ* = *ζ*_*c*_ := *Z*_*c*_*/*Λ^1*/*3^. The determination of *ζ*_*c*_ and *Z*\* are key goals of our analysis. The unpatterned region corresponds to *ζ < ζ*_*c*_ and the patterned region corresponds to *ζ > ζ*_*c*_. The solution is smaller in the unpatterned region, with the scaling *Y* = *W*_*L*_*/*Λ^1*/*3^, and larger in the patterned region, with the scaling *Y* = Λ^1*/*6^*W*_*R*_.

Using the scalings noted above, the leading-order scaled version of the system (6), (7) in the patterned state (*ζ > ζ*_*c*_) is

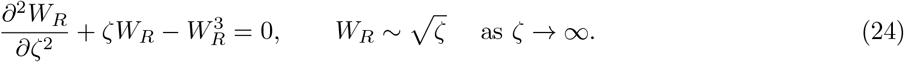

Similarly, the leading-order scaled version of the system (6), (7) in the unpatterned state (*ζ < ζ*_*c*_) is

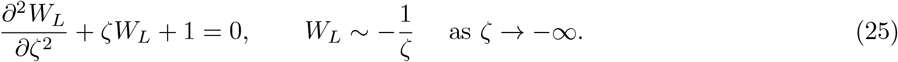

Since the nonlinear ODE in (24) is second order, it is more difficult to obtain an analytic solution within the patterned region than it was for Region B. However, we can solve (25) straightforwardly using the variation of parameters method to obtain

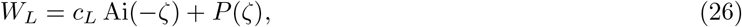

where the constant *c*_*L*_ can be obtained via matching with the solution in the patterned region, and where the function *P* (*ζ*) is defined as

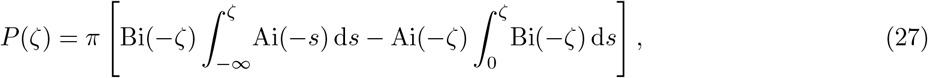

and is related to the Scorer functions [35, Sec. 9.12].

Given the relative scalings of *Y* in each asymptotic region, we can determine the position *ζ* = *ζ*_*c*_ by calculating when *W*_*R*_ = *O* (Λ^−1*/*2^). Similarly, the dynamic position of the bifurcation *Z*\* = Λ^1*/*3^*ζ*\* can be inferred through the relationship *Y* (*Z*\*) = 1, and hence *W*_*R*_(*ζ*\*) = Λ^−1*/*6^. Hence, we require knowledge of the decaying behaviour of the nonlinear ODE (24) as we move towards the unpatterned far field *ζ* → −∞. Linearising (24) around the far-field trivial solution and imposing decay results in the behaviour

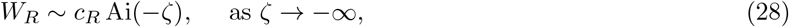

where *c*_*R*_ is an *O* (1) constant, which is straightforward to determine numerically if required. (Matching with the unpatterned region gives the relationship *c*_*L*_ = Λ^1*/*2^*c*_*R*_.)

Using the far-field behaviour of the Airy function, we find that *W*_*R*_ = *O* (Λ^−1*/*2^) for *ζ*_*c*_ (log Λ^3*/*4^)^2*/*3^ and that *W*_*R*_(*ζ*\*) = Λ^−1*/*6^ yields *ζ*\* (log Λ^1*/*4^)^2*/*3^, resulting in the following asymptotic result for the dynamic position of the bifurcation

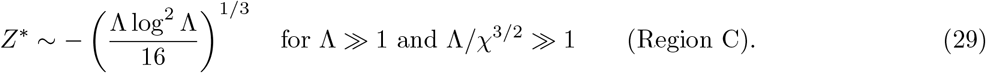

For a perfect pitchfork, as defined in (10), (11), this Region C limit is equivalent to Γ 1. In this case, the leading-order version of (10), (11) is exactly (24) with far-field decay as *ζ* → −∞. Noting that the perfect case has already been scaled into the appropriate form for the leading-order system here, we can infer that the extent of the transition region in this case will be *x* − *s*(*t*) = *O* ((*D/η*)^1*/*3^), and that the behaviour of *y* as *z* → −∞ will be *y* ∼ *c*_*R*_ Ai(−*z*) for some constant *c*_*R*_ = *O* (1).

#### 3.1.4 Summary

Our results suggest that the emergent dynamics of pattern-forming systems are determined in time and space by the regime in which the parameters lie (Fig. 1b-c). Furthermore, our results *quantify* how the location of a dynamic bifurcation is determined in each regime - either purely by a balance between the reaction terms for Region A, or otherwise for Regions B and C (Fig. 1b,c). In Regions B and C, diffusion and temporal variations in the parameters promote shifts in the location of the bifurcation that require an asymptotic analysis to quantify. While we give the specific (quantitative) asymptotic results in each section, the scalings for both pitchfork and transcritical bifurcations are similar and can be summarised as follows:

**Region A**. Reaction-dominated and quasi-steady patterning. In this case, Λ ≪ 1 and |*χ*| ≪1, and the reaction terms (iii)-(v) dominate in Eq. (6). The dynamical position of the bifurcation corresponds to its quasi-steady position, i.e. *Z*\* = 0 (Fig. 1b,c).

**Region B**. Critical slowing down. In this case |*χ*| ≫ 1 and Λ^2*/*3^*/* |*χ*| ≪ 1, and terms (ii)-(v) dominate in Eq. (6) – saturation, i.e. term (v), can be ignored in the unpatterned part of the domain, and base production, i.e. term (iii), can be ignored in the patterned p<u>art. The</u> dynamical position of the bifurcation lags behind its quasi-steady position, and scales as 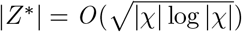 (Fig. 1b,c).

**Region C**. Diffusively enhanced, quasi-steady patterning. In this case Λ ≫ 1 and Λ^2*/*3^*/* |*χ*| ≫ 1, and terms (i), (iii)-(v) dominate in Eq. (6) – saturation can be ignored in the unpatterned part of the domain, and base production can be ignored in the patterned part. The dynamical position of the bifurcation is shifted via diffusion towards the unpatterned part of the domain, and scales as |*Z*\*| = *O* ((Λ log^2^ Λ)^1*/*3^) (Fig. 1b,c).

### 3.2 Spatio-temporal parameter oscillations for pitchfork bifurcations

In Section 3.1 above, we calculate the effective delay caused by travelling-wave-like spatio-temporal variations i.e. *monotonically* moving bifurcations, identifying and analysing three distinct parameter regions in space that demonstrate different types of behaviour. There are also important questions regarding robustness in biological systems involving oscillatory spatio-temporal variations i.e. *non-monotonically* moving bifurcations. As a key example of this - consider the situation where spatio-temporal variations in system coefficients would cause the system (at a fixed position in space) to move back and forth between the patterned and unpatterned regions predicted by a quasi-steady analysis (i.e. a local balance between reaction terms). How does the persistence or decay of the patterning depend on the spatio-temporal variation in the system? And can the delaying effects quantified in the previous section cause an effective robustness of the system to spatio-temporal variations over non-standard intermediate timescales, as seen in subsection 3.1? Quantitative answers to such questions are not covered by the analysis in subsection 3.1, which was restricted to monotonic bifurcation movement.

To determine whether non-monotonic moving bifurcations can have a significant effect on pattern persistence, we perform a preliminary numerical investigation of the effect of oscillatory variations in the system Eq. (1) (Fig. 1b). We find that there is an effective inertia associated with oscillating over the bifurcation, and so each system may indeed become stuck in the unpatterned or patterned state (Fig. 1b) for parameters oscillating quickly enough, even at large amplitude oscillations. This suggests that the bifurcations in each system can act as low-pass filters on parameter variations.

To understand why this happens and to quantify the phenomenon, we now investigate the system around a *fixed* position in space, in cases where the moving bifurcation turns around. Again, with the aim of balancing breadth and brevity, we consider the biologically relevant cases of patterned-unpatterned-patterned for pitchfork bifurcations (*n* = 3) in the main text, and unpatterned-patterned-unpatterned for transcritical bifurcations (*n* = 2) in Appendix B.2. Moreover, since we are specifically concerned with robustness to spatio-temporal variations, we restrict our analysis here to the ‘critical slowing down’ regime. This is the equivalent of Region B from the previous subsection, in which dynamics are slowed by passage through the bifurcation [31]. Specifically, this means we consider the scenario where diffusive effects are less important. However, we will retain the generality of the spatio-temporal variations by preserving their full nature and resisting the temptation to expand them locally in space and time.

#### 3.2.1 Deriving inner equation

Here, we consider (1) with *n* = 3, which produces a pitchfork bifurcation. Hence, we start with the system

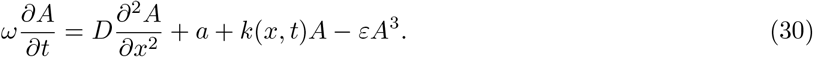

We are specifically interested in spatio-temporal oscillations of *k*(*x, t*) (i.e. cases where the bifurcation can turn around) and in understanding the intrinsic robustness of patterning due to this non-monotonically moving bifurcation. As such, we zoom into a region near the generic fixed point *x* = *x*\*, around the time *t* = *t*\* at which the bifurcation reaches the fixed point, defined through *k*(*x*\*, *t*\*) = 0. We consider the scenario where the point *x* = *x*\* transitions from patterned-unpatterned-patterned under a quasi-steady analysis (i.e. a local balance between reaction terms). Mathematically, this corresponds to *k*(*x*\*, *t*) transitioning from positive-negative-positive. When *k <* 0, there is a drive to de-pattern. Since we are specifically interested in understanding when the system demonstrates intrinsic robustness, we are interested in understanding when the system overcomes this forcing to de-pattern and patterning persists. Therefore, we investigate when the critical slowing down effect of Region B presented in Section 3.1 can overcome this forcing before *k* becomes positive again and the system returns to the locally patterned state. That is, quantifying when the system is robust to temporary incursions into regimes that would appear to be unpatterned from a quasi-steady analysis.

Since we are focusing on the critical slowing down scenario, diffusive effects can be neglected. As such, we can effectively consider *D* = 0 in (30) and in what follows. However, we retain *D* for the time being in order to understand when we can formally neglect it. As before, we treat *ω* and *ε* as small.

We are specifically interested in understanding when forcing transitions from patterned-unpatterned-patterned resist de-patterning and remain patterned, so the key terms in (30) are the left-hand side and the final two terms on the right-hand side. While we will formalize this intuitive dominant balance for the specific scenarios in which we are interested below, it is helpful to note that for this dominant balance all relevant asymptotic scalings will reduce (30) to an equation of the form

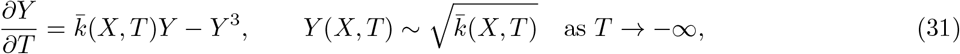

at leading-order, for appropriately defined independent variables *X, T*, dependent variable *Y* (*X, T*), and function 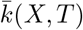, noting that 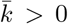 as *T* → −∞. It is straightforward to solve (31) by introducing *W* (*X, T*), defined through the transformation *Y* ^2^ = (∂*W/*∂*T*)*/*(2*W*), to turn (31) into

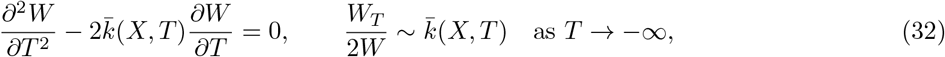

which can be solved in terms of 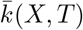, and transformed back into a solution for *Y* as follows:

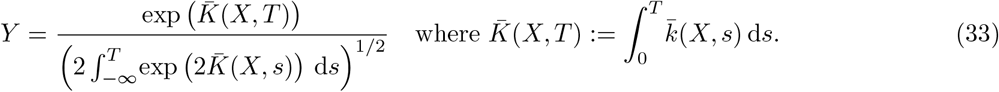

#### 3.2.2 Fixed away from the turning point

We first consider the case where *k*_*t*_(*x*\*, *t*\*) ≠ 0 i.e. the position at which the bifurcation turns around is away from the fixed point *x* = *x*\*. In this case, the appropriate inner equation scalings into the initially patterned region are

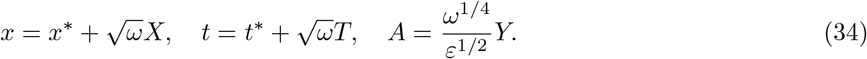

These scalings turn (30) into

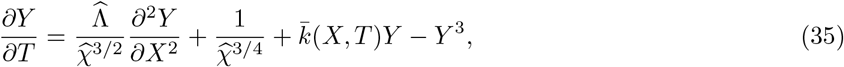

where we introduce the parameter groupings

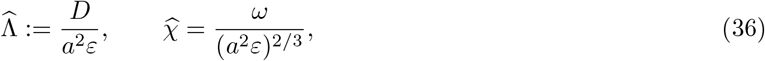

keeping similar notation as in (6b), and the function

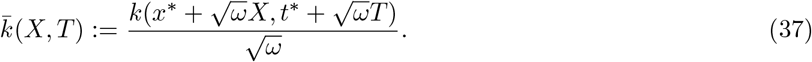

We note that 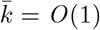 since *k*(*x*\*, *t*\*) = 0. Moreover, while it is possible to replace the right-hand side of (37) with its linearization for many differentiable functions *k*, we keep it in its general form since we are specifically interested in oscillations in *T*. That is, we are specifically interested in 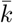 with turning points in *T*, and these will generally not be well-approximated by their linearizations.

The formal neglect of the diffusive and base production terms in (35) correspond to the limits 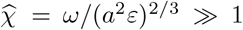 and 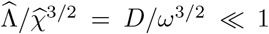, directly analogous to the limits that take us into Region B in the previous section. Moreover, this limit reduces (35) to the system (31), which is solved by (33). We can calculate when *Y* reaches a critical minimal value (specified by the particular problem being considered) via simple 1D root finding of the asymptotic solution (33) (see Fig. 4). While there will be additional asymptotic regions (analogous to their Region B versions) as *Y* becomes smaller for both imperfect and perfect pitchforks, we expect practical de-patterning to occur before these regions are reached so do not consider them further.

#### 3.2.3 Fixed at the turning point

We now consider the case where *k*_*t*_(*x*\*, *t*\*) = 0 and *k*_*tt*_(*x*\*, *t*\*) *>* 0 i.e. the bifurcation turns around at the fixed point *x* = *x*\*. In this case, the appropriate inner equation scalings into the initially patterned region are

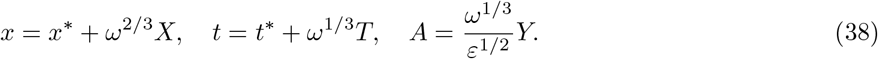

These scalings turn (30) into

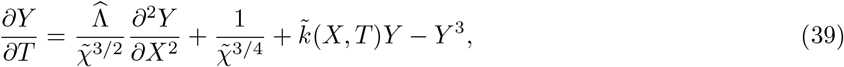

where 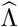 is defined in (36), and we introduce the parameter grouping

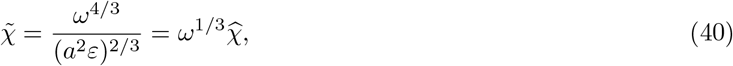

where 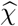 is defined in (36), as well as the function

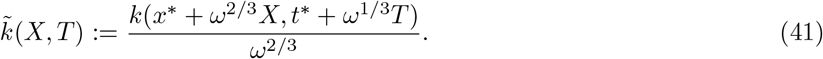

We note that 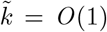 since *k*(*x*\*, *t*\*) = *k*_*t*_(*x*\*, *t*\*) = 0. Moreover, while it is possible to replace (41) with 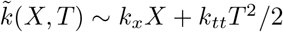 (with partial derivatives of *k* evaluated at (*x*\*, *t*\*)) for many differentiable functions *k*, we keep it in the form (41) for generality.

The formal neglect of the diffusive and base production terms in (39) correspond to the limits 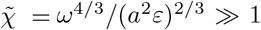 and 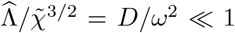. We note that both these constraints represent slightly stricter requirements on *ω* than their equivalents in the previous subsection, where we investigated a fixed position *away* from the turning point. Broadly, these constraints both require *ω* to be slightly larger than in the previous subsection for equivalent remaining parameters; this is because the constraints generally require 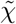 (and hence *ω*) to be large enough, and 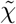 is a factor of *ω*^1*/*3^ smaller than 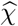, so needs to compensate. If these constraints are satisfied, this limit again reduces (39) to the system (31), which is solved by (33).

## 4 Biological example: Turing patterns during development

We apply our analytic results and perform simulations in the context of two paradigmatic biological pattern-forming systems with variations in pre-pattern morphogens. In each case, the morphogen variations affect parameters in the system gene-regulatory network [8]. In the main text here we consider a model for digit formation via activator-inhibitor Turing patterns [15], which is associated with a pitchfork bifurcation. In Appendix C, we consider a model for bacterial quorum sensing (QS) in *Vibrio fischeri*, which causes bioluminescence in the Hawaiian bobtail squid [36]. The bacterial quorum sensing system is associated with a transcritical bifurcation.

### 4.1 Mathematical model

We consider digit formation in the embryo, which has been modelled as a Turing system [15]. The kinetic parameters of the system are controlled by a spatio-temporally varying morphogen called fibroblast growth factor (Fgf) [15], which affects the self activation of the activator (Fig. 2a). For a minimalistic representation, and following a model in [15], we ignore the effects of other morphogens that are thought to affect the system. We expect our results to extend to more complicated systems that account for more morphogens and more complicated gene-regulatory networks [10, 17]. The system is associated with a pitchfork bifurcation.

**Figure 2:**
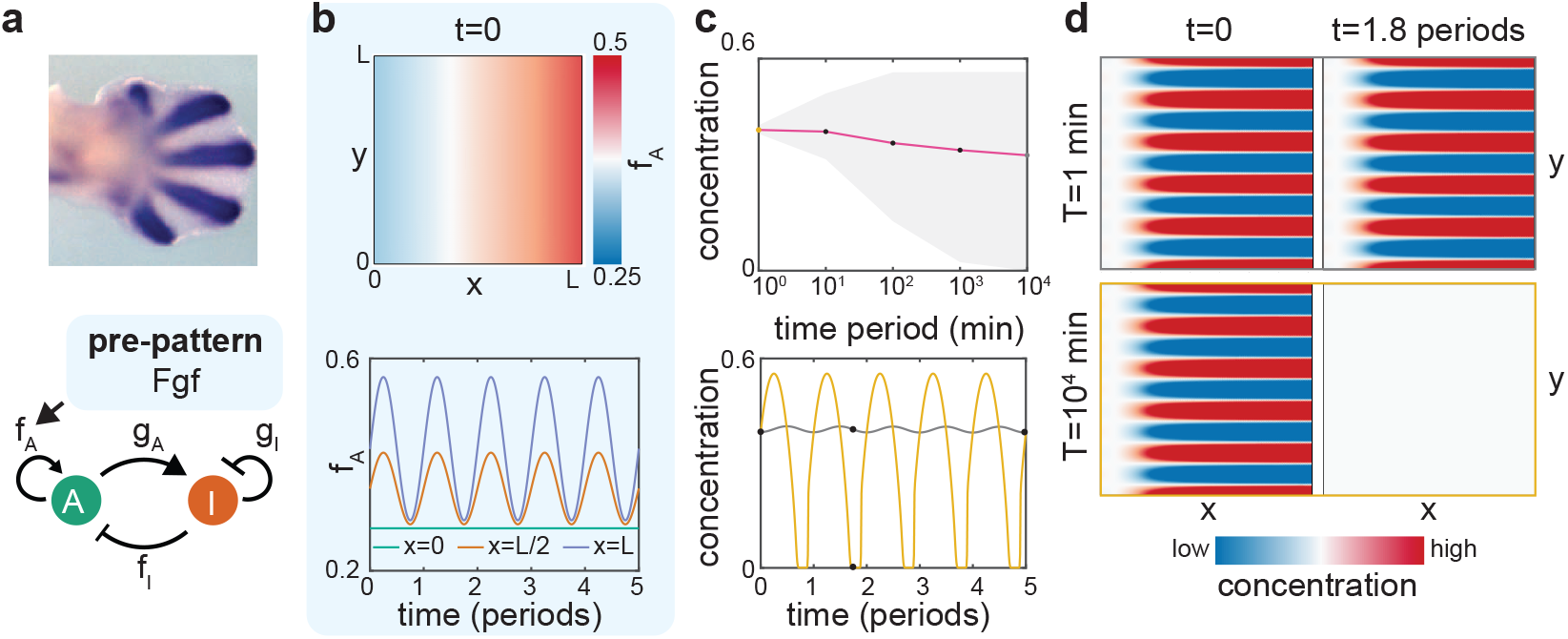
Effect of spatio-temporal morphogen variations on Turing patterns. a) We model Turing patterns during digit formation (top; reproduced from [15] with permission). The system’s pattern can be represented as an activator-inhibitor system, with the self-activation parameter *f*_*A*_ modulated by a morphogen called Fgf (bottom). b) The parameter *f*_*A*_ (min^−1^) at *t* = 0 (top) and throughout the spatio-temporal oscillations Eq. (44) (bottom; Movie S1). c) Top: Effect of time period 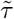 of spatio-temporal oscillations in Fgf on the mean concentration of the activator morphogen (line), and the range of concentration (grey area) during the oscillations. Bottom: Oscillations in the activator morphogen concentration for fast 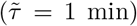 and slow 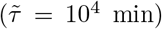 oscillations in Fgf (Movie S1). d) For oscillations that are filtered out, patterning remains the same (top). For oscillations that are not filtered out, patterning completely disappears (bottom). Images correspond to the black dots in panel c. Concentrations in the figure are non-dimensional and represent deviation from a base state at *x* = 0 [17].

The digit formation example we consider in this section is described by the equations

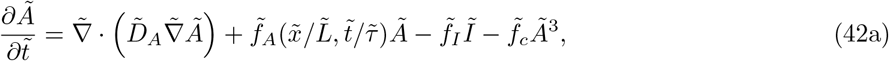

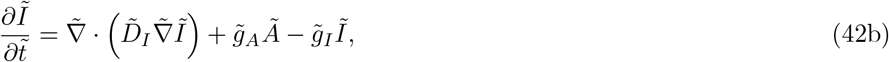

where the tildes denote dimensional quantities, and are supplemented with no-flux boundary conditions on the exterior of the domain

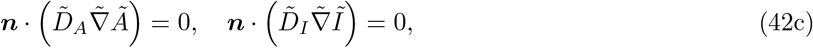

where *Ã* and *Ĩ* are the concentrations of the activator and inhibitor, respectively, 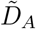 and 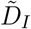 are diffusion coefficients, and 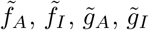 and 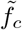 are kinetic parameters (Fig. 2a). To simulate spatio-temporal variations in Fgf [15], we let 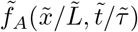 vary in space and time:

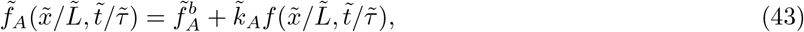

where 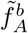 is the base self-activation of the activator, 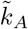 is a typical increase in the self-activation, and 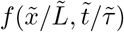 is a non-dimensional concentration of Fgf. Here, 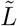 and 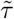 are typical lengthscales and timescales of the variation. For the numerical simulations we present in Fig. 2, we consider the system (42) in two spatial dimensions, but we restrict our analysis of the system to one spatial dimension.

We will generally use either

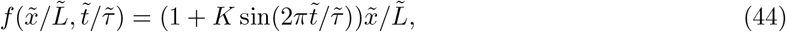

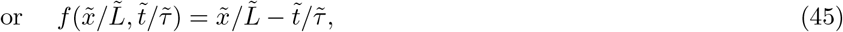

and we note that 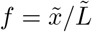 corresponds to the steady system studied in [15]. As an initial condition, we use the steady-state patterned solution for *K* = 0, with stripes oriented parallel to 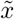, to test the effect of oscillations in Fgf on the patterns investigated in [15]. For simplicity, we use the same parameters as [15] for all kinetic parameters (Table 1). For the oscillation case (44), we set *K* = 0.9 to represent an oscillation in chemical concentration of around 90% over a timescale 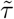; the aim is to find the critical 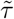 for which the oscillation is observed in the pattern-forming reactants.

**Table 1:** Inputs to model and outputs of the analysis of the Turing system.

| Symbol | Definition | Value and units | Reference |
| --- | --- | --- | --- |
| **Inputs** |  |  |  |
| $\tilde{L}$ | Length of domain in both dimensions | 717 $\mu\text{m}$ | [15] |
| $\tilde{f}_A^b$ | Base self-activation of activator | 0.28 $\text{min}^{-1}$ | [15] |
| $\tilde{f}_I$ | Inhibition of activator by inhibitor | 0.5 $\text{min}^{-1}$ | [15] |
| $\tilde{f}_c$ | Nonlinear saturation | 1 $\text{min}^{-1} \text{ nM}^{-2}$ | Defined to be consistent with [15] |
| $\tilde{g}_A$ | Activation of inhibitor by activator | 0.75 $\text{min}^{-1}$ | [15] |
| $\tilde{g}_I$ | Self-inhibition of inhibitor | 0.5 $\text{min}^{-1}$ | [15] |
| $\tilde{D}_A$ | Diffusion coefficient of activator | 70 $\mu\text{m}^2 \text{ min}^{-1}$ | [15] |
| $\tilde{D}_I$ | Diffusion coefficient of inhibitor | 875 $\mu\text{m}^2 \text{ min}^{-1}$ | [15] |
| $\tilde{k}_A$ | Average activation by Fgf at $x = \tilde{L}$ | 0.15 $\text{min}^{-1}$ | Chosen from [15], so that the system is patterned at $t = 0$ |
| $K$ | Size of oscillation in Fgf concentration | 0.9 | Chosen to test the effect of large oscillations in Fgf |
| $\tilde{\tau}$ | Time period of spatio-temporal variations | 1 day (this parameter also varied to find critical T) | [17] |
| **Outputs** |  |  |  |
| $\tilde{L}_d$ | Predicted extent of diffusive enhancement in comparison to $\tilde{L}$ | 7.2 % | |
| $\tilde{T}_{c,t}$ | Predicted critical timescale of variation of travelling gradient to switch between Region B and Region C | 5 hours 9 minutes | |

### 4.2 Nondimensional problem

We nondimensionalise the 1D version of (42) using

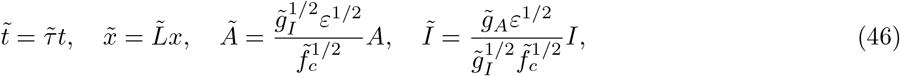

where the concentration scalings have been chosen to scale straightforwardly into the forthcoming weakly nonlinear analysis. The scalings (46) transform (42) into

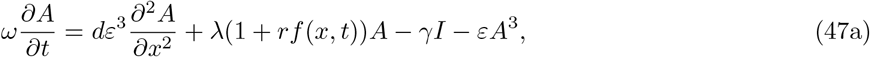

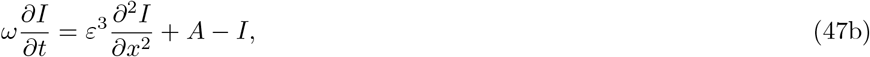

with

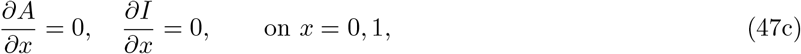

where we have introduced the dimensionless parameters

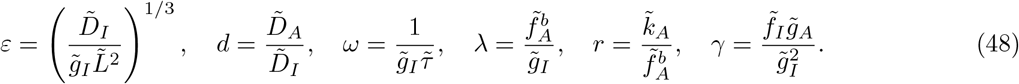

We will proceed by reducing this system to its inner normal form, via a systematic asymptotic analysis in which we formally exploit the small parameters *ε, ω* ≪ 1.

### 4.3 Reduction to canonical form

#### 4.3.1 Travelling-wave-like parameter variations

The relevant inner scalings for this problem involve two spatial lengthscales. These consist of a short scale *X* that characterises the small wavelength of patterns, and an intermediate scale *σ* that characterises the (more) slowly varying amplitude of the patterns. Mathematically, these are defined as

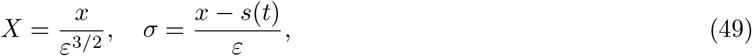

where the *O* (*ε*^3*/*2^) scaling for *X* is determined by balancing the diffusive terms with the linear reaction terms, and the *O* (*ε*) scaling for *σ* is determined by balancing the linear reaction terms with the nonlinear saturation terms. The scalings (49) transform the governing equations (47) into the system

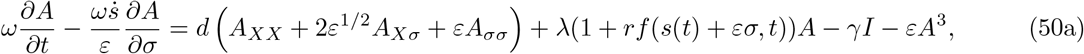

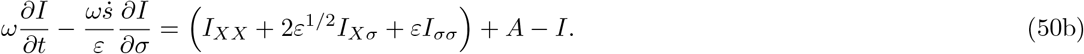

We proceed by considering an asymptotic expansion in powers of *ε*^1*/*2^

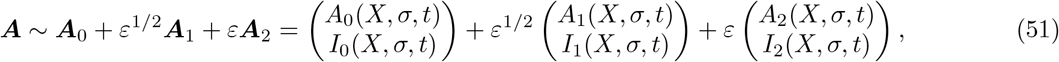

where the leading-order solution can be obtained through a straightforward linear analysis of the system. The parameter *ω* ≪ 1; it is convenient for our analysis to treat *ω* = *O* (*ε*^2^), which we shall see represents an important distinguished limit of the system.

At *O* (1), we obtain the leading-order system

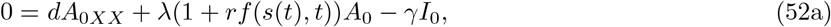

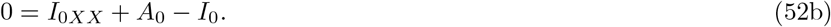

Since we are analysing the system near the Turing bifurcation, we seek the degenerate solution near *x* = *s*(*t*), the position of which is to be determined.

Since *σ* and *t* effectively act as parameters in (52), analysis of the linear system (52) proceeds in a similar manner to a classic linear stability analysis for Turing patterning in a homogeneous environment. To this end, the position of the moving bifurcation *x* = *s*(*t*) is defined implicitly through the relationship

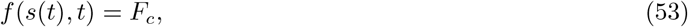

for constant *F*_*c*_ to be determined as part of our analysis. The value that *F*_*c*_ takes at the onset of bifurcation will be the same as the value it would take in the scenario of a homogeneous environment (as recently shown more broadly in [11] for spatially varying environments).

Given (53), we write (52) as

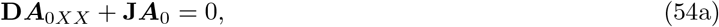

where

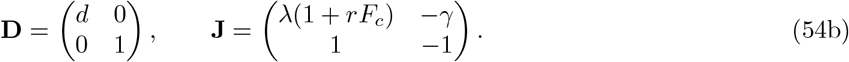

The onset of the Turing bifurcation is then given by (see, for example, [37]) the criteria

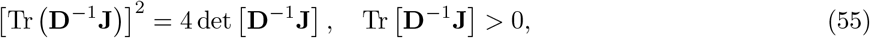

which combine to yield the following result for *F*_*c*_:

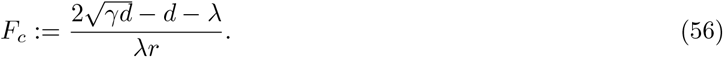

The result (56) allows us to write **J** in (54b) as

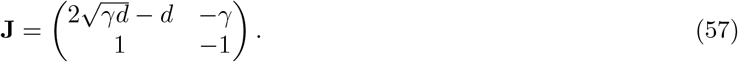

The remaining task at this order is to solve the system (54). In the standard manner, we look for a real solution proportional to 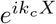, where *k*_*c*_ is the (as-of-yet unknown) wavelength of the patterned solution at the onset of bifurcation. Using this form of the solution, we see that 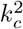 corresponds to the *repeated* eigenvalues of **D**^−1^**J** at the onset of bifurcation, and hence that

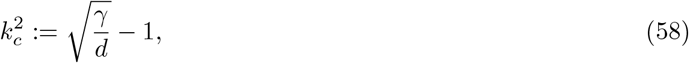

noting that 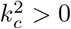 for Turing patterning to be possible. Hence, the system (54) is solved by

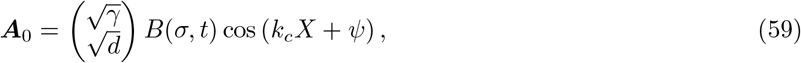

where 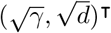 is the right eigenvector of **D**^−1^**J**, and the slowly varying amplitude *B* satisfies some governing equation that must be derived at higher asymptotic orders via our weakly nonlinear analysis. The derivation of this governing equation is the goal of our remaining analysis. Finally, we note that *ψ* is some phase shift that would have to be obtained via matching with a nonlinear outer solution. We note that knowledge of *ψ* is not required to capture the emergent amplitude nor frequency of patterning, since it only represents a translational shift in the patterning.

The *O* (*ε*^1*/*2^) terms in (50) yield

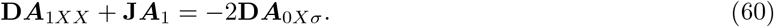

Since the right-hand side of (60) is orthogonal to the left-eigenvector of the linear operator, we do not have enough information to impose a meaningful solvability condition at this order. Hence we must proceed to the next order to yield a governing equation for *B*. Nevertheless, we must still solve (60) in terms of ***A***_0_, since the solvability condition we will derive at *O* (*ε*) requires information about ***A***_1_.

Using the solution (59), we see that the particular integral of (60) will be proportional to sin(*k*_*c*_*X* + *ψ*). Looking for a particular integral in this form, we find that

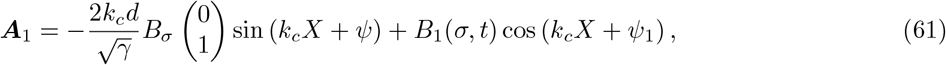

where the first term on the right-hand side is the particular integral of (60), and the second term on the right-hand side is the complementary function. *B*_1_ is an undetermined function of *σ* and *t*, and *ψ*_1_ is a constant that would be determined through matching at higher orders. We will not calculate either of these, since neither are required for our overarching goal: to derive a governing equation for *B*. This remains our objective, and to achieve this we proceed to the next asymptotic order.

The *O* (*ε*) terms in (50) yield

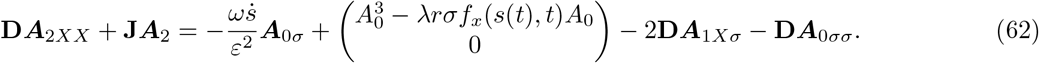

We can obtain the governing equation we seek for *B* by imposing a solvability condition on (62). To do this, we left-multiply the system (62) by its left-eigenvector (adjoint) solution 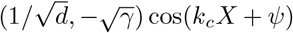 and integrate over a single period in *X* e.g. from *X* = 0 to 2*π/k*_*c*_. This procedure yields the following governing equation for *B*:

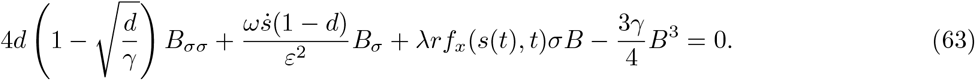

Finally, we note that we can rescale (63) into the normal form we give in (10) through the scalings

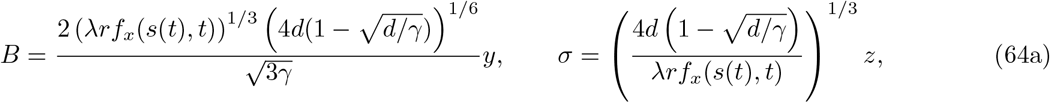

with

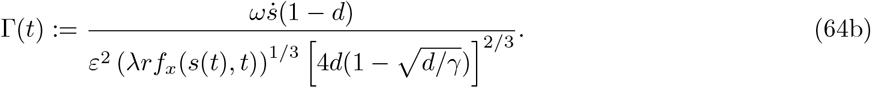

#### 4.3.2 Spatio-temporal parameter oscillations

We now consider the case of spatio-temporal parameter oscillations i.e. where the moving bifurcation can turn around. This is of particular interest in determining whether patterning persists dynamically even when an equilibrium analysis might suggest the repeated emergence and disappearance of a bifurcation at the edge of the domain. To analyse and quantify this effect, we scale into the region close to a generic fixed point *x* = *x*\*, around the time *t* = *t*\* at which the bifurcation reaches this point. That is, *t*\* is defined implicitly through *f* (*x*\*, *t*\*) = *F*_*c*_. We will later use this result to consider the potential persistence of a pattern-forming bifurcation at the edge of the domain.

We are specifically interested in the critical slowing down region where delayed effects dominate. We therefore use the new length, time, and concentration scalings

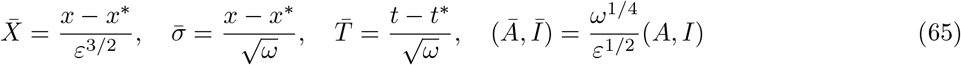

motivated by the fast wavelength scaling for *X* in (49), and the time, space, and concentration scalings in (34).

After applying the scalings (65), the resulting analysis is very similar to that in the previous subsection, so we omit it for brevity. This analysis yields the general leading-order solution

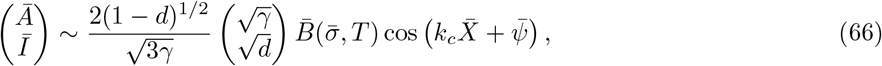

where the constants premultiplying this solution are added for notional convenience in presenting the following amplitude equation satisfied by the slowly varying amplitude 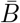:

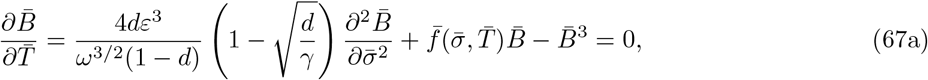

where

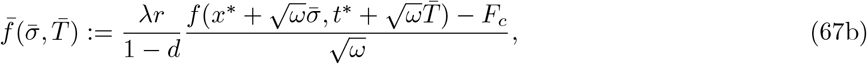

noting that *F*_*c*_ is defined in (56). Then, we see that if *ε* ≪ *ω*^1*/*2^, the system (67) reduces to (31), which has solution (33). Hence, for a specified function *f*, we can calculate the value of 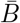 through the general solution (33). Moreover, for finite domains *x* ∈ (0, 1), if we apply this analysis at the boundary *x* = 1,^2^ we can calculate the critical oscillation period for which patterns persist (i.e. remain above a specified critical value) via simple 1D root finding (see Fig. 4).

## 5 Biological examples: Results and implications

So far, we have analytically studied two biological pattern-forming systems: digit formation in the embryo (Section 4) and bacterial quorum sensing (Appendix C). Our modelling assumptions and the governing equations for each system are given at the beginning of Section 4 (Turing patterns) and Appendix C (quorum sensing); corresponding schematic diagrams are given in Fig. 2a and Fig. 3a, respectively. In both systems, we have derived results that quantify the dynamical and spatial effects of spatio-temporal pre-pattern morphogen variations. In this section, we link the analytical results in Section 4 and Appendix C to the robustness of biological pattern formation in pre-pattern morphogen variations. We also supplement the analytical results with the results of 2D simulations of the governing equations of each system in the finite-element computational software COMSOL Multiphysics.

**Figure 3:**
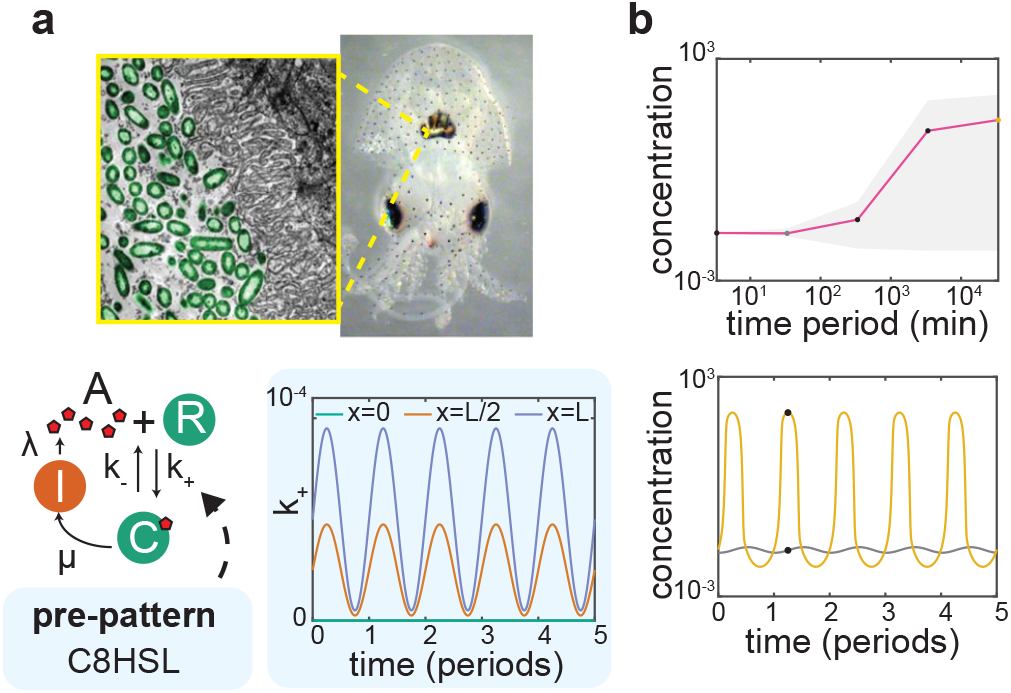
Effect of spatio-temporal morphogen variations on bacterial quorum sensing in a small biofilm or cell population. a) We model quorum sensing in *V. fischeri*, which causes bioluminescence in the Hawaiian bobtail squid (top; images adapted from [38], with permission). We model the LuxR system, in which an autoinducer, 3OC6HSL, promotes its own synthesis by binding with a protein, LuxR, to form a transcription factor (bottom left). The binding parameter *k*_+_ is modulated by competitive binding with a second autoinducer, C8HSL. Bottom right: the parameter *k*_+_ (nM^−1^s^−1^) throughout the spatio-temporal oscillations Eq. (C.4) (see also Movie S2). b) Top: Effect of the time period 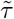 of the spatio-temporal oscillations in C8HSL on the mean 3OC6HSL concentration (line), and the range of 3OC6HSL concentration (grey area) during the oscillations. Bottom: Oscillations in 3OC6HSL concentration, for fast 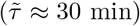 and slow 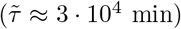 oscillations in C8HSL. For oscillations that are filtered out, the system remains at low 3OC6HSL concentration. For oscillations that are not filtered out, the cell population fills with 3OC6HSL during the oscillation (see Movie S2). Concentrations in the figure are scaled by the quorum sensing activation threshold of 5 nM [12].

We begin by assuming for simplicity that pre-pattern morphogen variations act like a ‘travelling wave’ of the form Eq. (45). Our analysis in Section 4 and Appendix C shows that in this scenario, the governing equations of both biological systems reduce to the weakly nonlinear normal forms Eq. (6a) or Eq. (10). The pre-pattern morphogen variations are then encapsulated in two parameters, Λ(*t*) and *χ*(*t*), defined through the ratio Γ(*t*) := *χ/*Λ^2*/*3^ in (64b) for the digit formation application and in (C.27) for the quorum sensing application. Once Λ(*t*) and *χ*(*t*) are calculated, the dynamics of the system are classified through the general parameter space shown in Fig. 1C. The parameter space contains three regions in which we predict qualitatively different solution behaviour (Section 3); we have named them Region A (reaction-dominated; quasi-steady), Region B (critical slowing down), and Region C (diffusively enhanced; quasi-steady). These results demonstrate that both of the paradigmatic pattern-forming systems considered here are subject to universal regimes that we have identified, which is a consequence of their bifurcation structures.

To understand the biological implications of the solution regimes in Fig. 1C, we now apply our analytical results to identify which regimes are most relevant to each biological system, and to quantify the dynamics of patterning in those regimes. First, we focus on the effects of pre-pattern morphogen variations that occur over physiological timescales. We use physiologically relevant values of the parameters in both systems (Tables 1, 2), with the timescale of variation 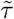 in the pre-pattern morphogen ‘travelling wave’ (Eq. 45) corresponding to the relevant growth timescale; this is in line with experimental evidence that suggests physiological changes to patterning in each system are induced by growth [15, 17, 39]. Our analysis predicts that for such parameters, each system sits in Region C of parameter space (diffusively enhanced patterning; see Fig. 1c). In Region C, dynamics are quasi-steady, so we would expect both systems to respond promptly to temporal changes in morphogens that occur over physiological timescales. Also in Region C, patterning is ‘enhanced’ through diffusion so that areas ahead of the pre-pattern morphogen ‘travelling wave’ become patterned (a precise summary of this effect is given in Section 3). We predict significant quantitative differences in the spatial extent of patterning between the two systems because of this effect. In the Turing system, the diffusive enhancement to patterning is less than 10% of the size of the domain (see Table 1) – patterning would be expected to be controlled relatively locally by the concentration of the pre-pattern morphogen Fgf. This prediction is in line with recent analyses of Turing systems in steady pre-pattern morphogen gradients, in which patterning was found to be controlled relatively locally in space [11]. By contrast, our analysis predicts that diffusive enhancement to patterning is much stronger in the bacterial quorum sensing system. We find that diffusive enhancement causes the patterned (or QS-activated) region to be larger than the size of the population (Table 2) – any local activation in QS caused by changes over a growth timescale would be expected to cause the entire population to activate. This would benefit the population by ensuring all cells commit together to a multicellular program of gene expression, as was recently found in fluid flows [12]. These results suggest that biological pattern-forming systems can tune via diffusion the extent of patterning.

**Table 2:**
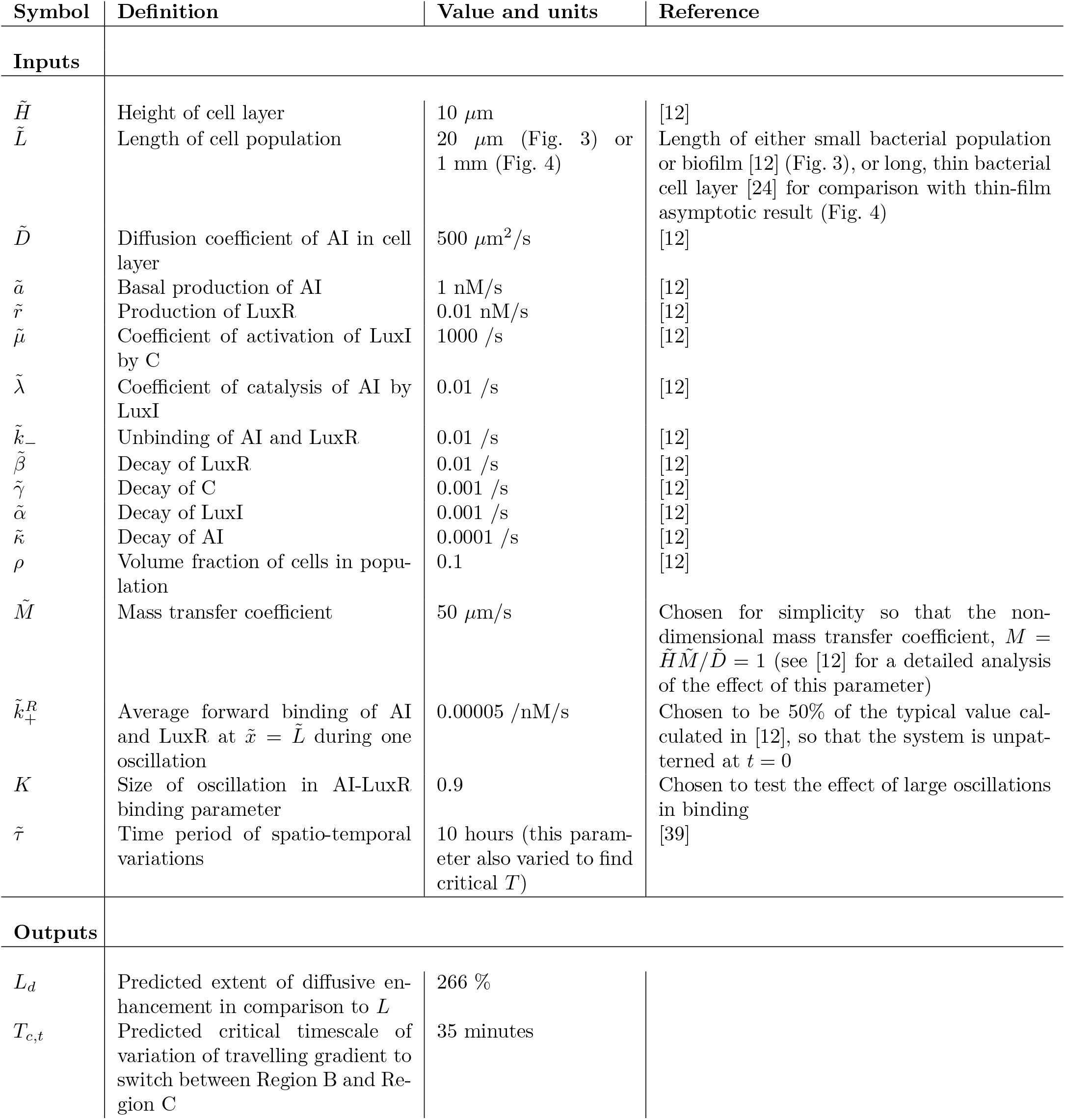
Inputs to the model and outputs of the analysis of the quorum sensing system.

| Symbol | Definition | Value and units | Reference |
| --- | --- | --- | --- |
| **Inputs** |  |  |  |
| $\tilde{H}$ | Height of cell layer | 10 $\mu\text{m}$ | [12] |
| $\tilde{L}$ | Length of cell population | 20 $\mu\text{m}$ (Fig. 3) or<br>1 mm (Fig. 4) | Length of either small bacterial population or biofilm [12] (Fig. 3), or long, thin bacterial cell layer [24] for comparison with thin-film asymptotic result (Fig. 4) |
| $\tilde{D}$ | Diffusion coefficient of AI in cell layer | 500 $\mu\text{m}^2/\text{s}$ | [12] |
| $\tilde{a}$ | Basal production of AI | 1 nM/s | [12] |
| $\tilde{r}$ | Production of LuxR | 0.01 nM/s | [12] |
| $\tilde{\mu}$ | Coefficient of activation of LuxI by C | 1000 /s | [12] |
| $\tilde{\lambda}$ | Coefficient of catalysis of AI by LuxI | 0.01 /s | [12] |
| $\tilde{k}_-$ | Unbinding of AI and LuxR | 0.01 /s | [12] |
| $\tilde{\beta}$ | Decay of LuxR | 0.01 /s | [12] |
| $\tilde{\gamma}$ | Decay of C | 0.001 /s | [12] |
| $\tilde{\alpha}$ | Decay of LuxI | 0.001 /s | [12] |
| $\tilde{\kappa}$ | Decay of AI | 0.0001 /s | [12] |
| $\rho$ | Volume fraction of cells in population | 0.1 | [12] |
| $\tilde{M}$ | Mass transfer coefficient | 50 $\mu\text{m}/\text{s}$ | Chosen for simplicity so that the non-dimensional mass transfer coefficient, $M = \tilde{H}\tilde{M}/\tilde{D} = 1$ (see [12] for a detailed analysis of the effect of this parameter) |
| $\tilde{k}_+^R$ | Average forward binding of AI and LuxR at $\tilde{x} = \tilde{L}$ during one oscillation | 0.00005 /nM/s | Chosen to be 50% of the typical value calculated in [12], so that the system is unpatterned at $t = 0$ |
| $K$ | Size of oscillation in AI-LuxR binding parameter | 0.9 | Chosen to test the effect of large oscillations in binding |
| $\tilde{\tau}$ | Time period of spatio-temporal variations | 10 hours (this parameter also varied to find critical $T$) | [39] |
| **Outputs** |  |  |  |
| $L_d$ | Predicted extent of diffusive enhancement in comparison to $L$ | 266 % | |
| $T_{c,t}$ | Predicted critical timescale of variation of travelling gradient to switch between Region B and Region C | 35 minutes | |

With the aim of understanding robustness, we next investigate the effect of spatio-temporal morphogen variations that occur faster than growth. Interestingly, by reducing the timescale of variation 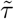 in the prepattern morphogen ‘travelling wave’ (Eq. 45), we find that each system transitions to Region B (critical slowing down) of parameter space at a critical value of 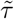 (Tables 1, 2). In Region B, a bifurcation-associated emergent timescale slows the system dynamics (see Section 3) – this suggests that both systems may be slow to respond to pre-pattern morphogen variations that happen over timescales that are not physiological. To this investigate this further, we now consider oscillatory pre-pattern morphogen variations of the form Eq. (44) in each system (see Movies S1, S2); in a quasi-steady system, we would expect to see period cycles in which patterning switches on and off. However, our analytical results show that in this scenario, for both pitchfork and transcritical bifurcations, there is a critical oscillation timescale – oscillations that occur faster than this timescale are predicted to be ‘filtered out’ because of the critical slowing down (see Section 4 and Appendix C). Our simulations confirm this finding: for pre-pattern morphogen oscillations that occur faster than a critical timescale, we find that both biological systems remain stuck in the initial patterned or unpatterned state (Fig. 2c,d; Fig. 3b; Movies S1, S2). We calculate the critical timescale to be around 3-10 hours, depending on the kinetic parameters in each system (Fig. 2c; Fig. 3b; Fig. 4) – surprisingly, this is a few hours faster than the timescale of growth in both systems (Tables 1, 2). Therefore, we expect changes in the pre-pattern morphogen concentration to be ignored if they occur much faster than a growth timescale, allowing the system to avoid repeated cycles of complete removal of patterning (Fig. 2d, Fig. 3b; Movies S1, S2). Remarkably, this suggests that the gene-regulatory parameters in each biological system are tuned such that each system is robust to non-physiological variations in pre-pattern morphogens.

**Figure 4:**
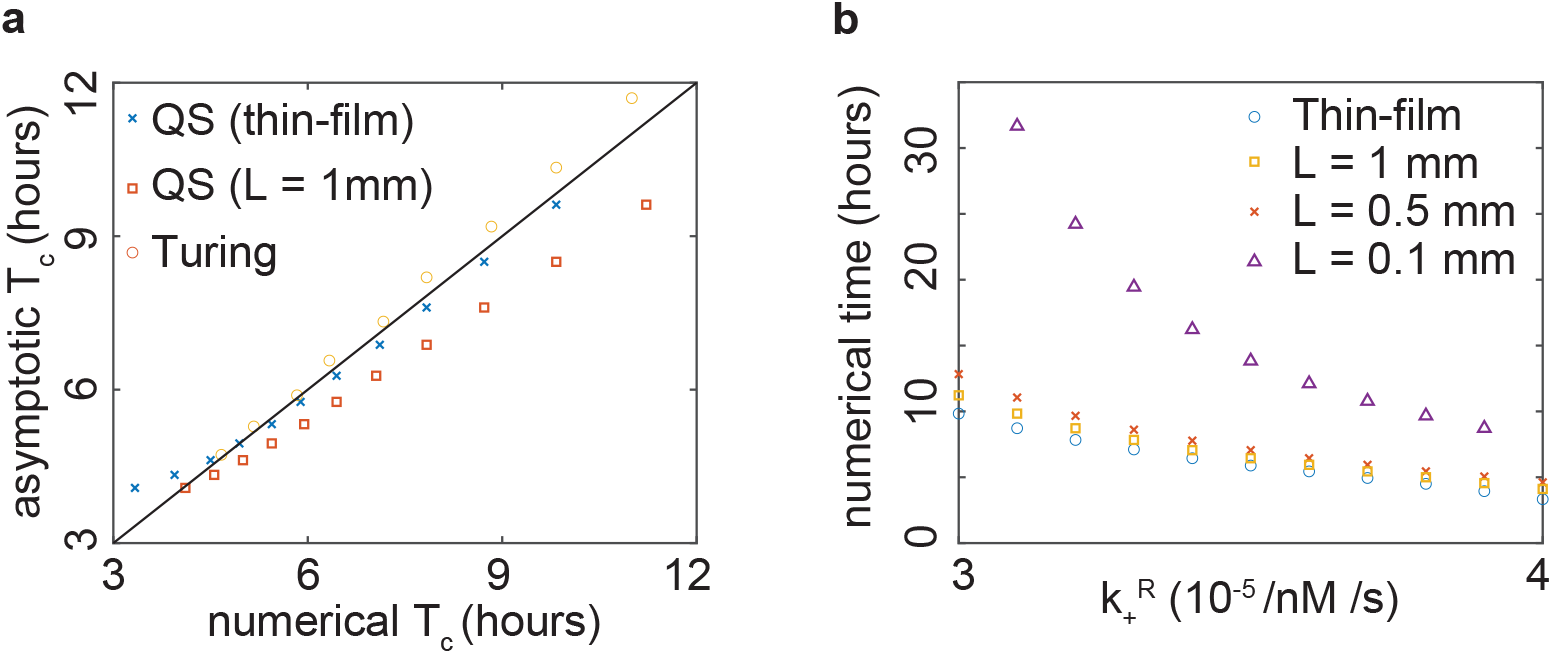
(a) Comparison of asymptotic and numerical values of critical time period for spatio-temporal oscillations. In the Turing system, we use the patterned state as the initial condition, to test how robust patterning is to spatio-temporal variations in the pre-pattern morphogen. We calculate a critical oscillation time from the simulations by finding the smallest oscillation period such that the concentration *A* falls below a critical value, equal to the wavelength of the predicted fastest growing mode at the onset of instability. This provides a natural criterion for the switch on or off of patterning that is independent of initial conditions. We compare to the corresponding asymptotic critical timescale with the same critical value, calculated via root finding of the asymptotic solution Eq. (33), which solves the ODE (31) to which (67) reduces in the limit *ε ω*^1*/*2^. In this figure, 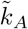 takes values between 0.15 /min and 0.8 /min. In the bacterial quorum sensing system, we use the unpatterned state as the initial condition, to test the robustness of the decision to implement a gene expression program in response to quorum sensing activation. In this system, diffusion is more important, so Eq. (31), which was essentially derived in the thin-film limit, requires a long, thin population to be quantitatively accurate. To illustrate its accuracy, we perform simulations for *L* = 1 mm. We also perform thin-film simulations in which 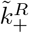 depends only on *t*, by setting a spatially uniform sinusoidal oscillation in Eq. (C.3). In this figure 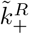 has values between 3 × 10^−5^ /nM/s and 4 × 10^−5^ /nM/s, so that the system remains in the unpatterned regime at *t* = 0 in the thin-film limit. We calculate a critical oscillation time from the simulations by finding the smallest oscillation period such that the concentration *A* rises above its critical value given in Table 2. We compare to the corresponding critical timescale (with the same critical value) calculated via root finding of Eq. (B.15), which solves (B.14), to which (C.29) reduces in the limits *ω* ≪ *δ*^4*/*3^ and *ω* ≪ *ε*. (b) The simulations of the bacterial quorum sensing system collapse onto the results of the thin-film simulations as *L* is increased to around 1 mm. For longer populations, the critical oscillation times shown for the quorum sensing system in panel a (crosses) is expected to be a quantitatively accurate approximation to the critical oscillation time. In all simulations in both panels, we allow the system to settle into periodic temporal oscillations before extracting results.

## 6 Discussion and conclusion

This study concerns biological pattern-forming systems with spatio-temporal variations in pre-pattern morphogens; we assume that the pre-pattern morphogens affect the kinetics of the underlying gene-regulatory network. Mathematically, this corresponds to variations in the parameters of the governing reaction-diffusion equations; the onset (or offset) of pattern formation is induced when variations cause the system to transition through a bifurcation. Our analysis of such systems demonstrates how the dynamics of biological pattern formation in pre-pattern morphogen variations can be classified and quantified in terms of universal solution regimes that we have identified. Each solution regime displays qualitatively different dynamics – the regime in which a biological system sits depends on the gene-regulatory kinetics, and on the timescale of the pre-pattern morphogen variations.

This presents the intriguing possibility that, with the appropriate gene-regulatory kinetics, a biological system could take advantage of different solution regimes in different situations. Surprisingly, our results indeed suggest that in both of the biological systems that we have studied, the system transitions into the ‘critical slowing down’ regime if variations in pre-pattern morphogens become much faster than growth. In this ‘critical slowing down’ regime, the onset (or offset) of pattern formation occurs over a new, longer, timescale that emerges because of the presence of a bifurcation in the system. We predict that this allows the biological systems to ‘filter out’ non-physiogical oscillations in pre-pattern morphogens. Overall, our results suggest that bifurcations in reaction-diffusion systems, which are associated with the onset of patterning, can also provide inherent robustness to biological systems.

In principle, any pattern-forming system that undergoes a transcritical or supercritical pitchfork bifurcation can be analysed using our asymptotic framework and plotted on the parameter space that our analysis has produced (Fig. 1c). We expect that our results can also be extended to wider classes of systems, such as those with Hopf bifurcations in their underlying governing equations. Our general framework complements recent work on pattern formation in various systems of equations with spatio-temporally varying parameters (e.g. [11, 40–47]) and on the effect of critical slowing down in a range of contexts [48, 49]. Furthermore, our classification of the dynamics of gene-regulatory network architectures via their low-dimensional mathematical structure (i.e. bifurcations) complements recent work on dimensionality reduction of systems without spatio-temporal heterogeneity [29, 50, 51] and systems transitioning from a dynamic to a static regime [22].

We have applied our results to two specific biological pattern-forming systems. In modelling these systems we have performed significant simplifications for clarity and generality. In particular, we have not modelled the effect of stochastic noise in pre-pattern morphogen concentrations, which we expect to have effects that are not captured by our analysis [52, 53]. Furthermore, our models are effective macroscopic representations of microscopic processes, and the process of coarse-graining the microscopic dynamics to obtain effective macroscopic dynamics is often not trivial [54].

To conclude, we have presented a general framework that classifies and quantifies the dynamic response of pattern-forming systems to spatio-temporal variations in their parameters. We have applied our framework to simple models of two biological pattern-forming systems, each with variations in a pre-pattern morphogen that affects kinetic parameters: digit formation via Turing patterns, and bacterial quorum sensing. Our theory predicts that both systems filter out spatio-temporal morphogen variations that occur much faster than growth. We demonstrate that the type of bifurcation in the system, which is determined by the gene-regulatory network, controls emergent patterning dynamics and structure. Predictions such as these are testable in newly developed systems that allow spatio-temporal control over gene-expression and the external environment, such as synthetic model organisms [55], organoids [56] and microfluidic devices [24]. Owing to the generality of the canonical equations that we have analysed, our theoretical framework is extendable to a wide class of pattern-forming systems.

## Acknowledgements

We thank Zena Hadjivasiliou and Jake Cornwall Scoones for helpful discussions, and Eric Stabb for assistance with Fig. 3a. MPD is supported by the UK Engineering and Physical Sciences Research Council [Grant No. EP/W032317/1]. PP is supported by a UKRI Future Leaders Fellowship [MR/V022385/1].

## A Detailed example of critical slowing down caused by a transcritical bifurcation

Consider the (dimensional) ODE system

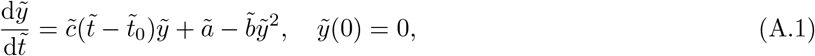

where 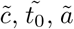, *ã*, and 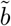 are positive constants. Here, 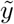 represents the concentration of some chemical species whose reaction rate is governed by a self-activation/self-repression term with time-varying strength 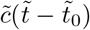, a base rate of production *ã*, and a saturation (self-repressing) effect with strength 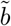. We non-dimensionalize using

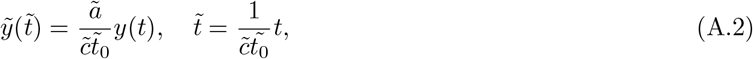

to obtain

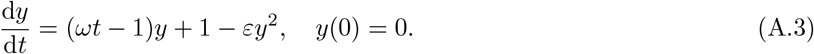

where

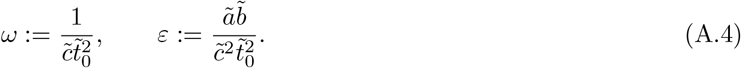

We are specifically interested in the asymptotic limit *ω, ε* ≪ 1. This corresponds to a fast base production (over *t* = *O* (1)) with a weak saturation effect, and a fast self-repression (over *t* = *O* (1)) that slowly varies (over *t* = *O* (1*/ω*)) until it becomes a self-activation. The point at which this switch between repression and activation occurs (*t* = *t*_*c*_ := 1*/ω*) is an effective imperfect transcritical bifurcation in the ODE. While the system (A.3) may appear to straightforwardly yield quasi-steady (local equilibrium) solutions, we will show that the crossing of the bifurcation can lead to delayed effects over perhaps unexpected timescales.

As noted above, there are clear ‘fast’ and ‘slow’ timescales in this problem. The fast timescale is *t* = *O* (1), which represents the fast transition from the initial condition to the quasi-steady solution. This is also the restorative timescale for any perturbations to the problem e.g. due to a noisy environment. Over this timescale, the leading-order solution is given by

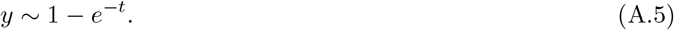

The slow timescale is *t* = *O* (1*/ω*), which represents the slow variation in the time-dependent coefficient. Over this timescale, it is helpful to explicitly introduce the slow timescale *T* = *ωt* = *O* (1), and to note that in this notation the fast timescale corresponds to *T* = *O* (*ω*). Transforming to the slow timescale *T*, the ODE (A.3) becomes

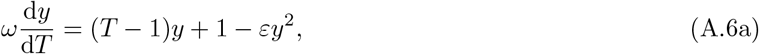

with appropriate matching condition

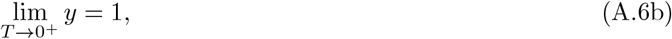

obtained through asymptotic matching with (A.5). The system (A.6) represents the apparent quasi-steady regime, and the corresponding leading-order solution is

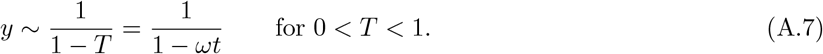

The apparent singularity in this solution occurs at the effective bifurcation point of the system: *T* = *T*_*c*_ := *ωt*_*c*_ = 1, where the slowly varying coefficient crosses zero. The apparent singularity explicitly highlights that there may be additional interesting behaviour as 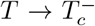. We will show that this interesting behaviour can lead to delays in the solution that occur over intermediate timescales in the problem of 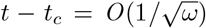, which is *slower* than the fast timescale of *t* = *O* (1) but *faster* than the slow timescale of *t* = *O* (1*/ω*).

To justify the appropriate scaling for the intermediate slowing timescale, it is helpful to jump forward in time slightly and understand the quasi-steady solution once *y* has become very large and saturation effects have kicked in to balance self-activation. The appropriate balance for this is between the (*T* − 1)*y* and *εy*^2^ terms in the ODE in (A.6a), which yields a solution of

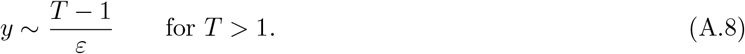

To motivate the appropriate intermediate timescale, we identify when the solutions (A.7) and (A.8) are of the same asymptotic order near the bifurcation point (*T* = *T*_*c*_ := 1). This occurs when 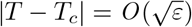, and hence when 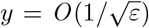. Therefore, we introduce new intermediate scalings, starting with an intermediate timescale *τ* = *O* (1) where 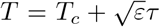. Note that this is equivalent to 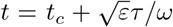. We also introduce the intermediate dependent variable *Y* (*τ*), defined through 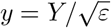. These scalings convert the ODE (A.6a) into

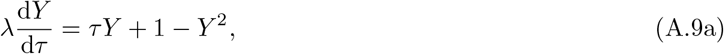

where 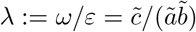, with appropriate matching condition

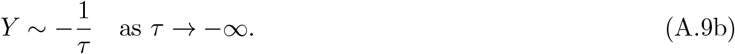

The system (A.9) represents the full intermediate dynamics of the system, and its behaviour is completely controlled by the single parameter grouping *λ*. Since we made no additional assumptions on the relative sizes of *ω* and *ε*, the parameter grouping *λ* can be small or large. While we can write down the exact solution to (A.9) in terms of hypergeometric functions, its exact form is not particularly informative. It is more instructive to examine the cases of small and large *λ* asymptotically, as this will allow us to understand the emergence of the delaying effect over unexpected timescales due to dynamic crossing of the bifurcation. However, even before doing this systematically, we can note that the parameter *λ* encodes how important the reaction rate *λ*d*Y/*d*τ* is directly from the form of the intermediate ODE (A.9). As such, we would broadly expect dynamic crossing effects to be important for large *λ* and unimportant for small *λ*.

Specifically, in the limit of *λ* ≪ 1 (equivalently *ω* ≪ *ε*), the system will remain quasi-steady throughout the intermediate timescale. In this case we can explicitly write down the asymptotic solution

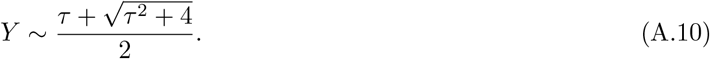

Hence, for small *λ* we see that the intermediate solution (A.10) provides a smoothing between the two outer solutions (A.7) and (A.8), but does not contribute any dynamic effects to the system. A helpful metric to gauge the level of delay in the system due to the bifurcation crossing is the delay between the dynamic solution and what would be expected in the quasi-steady system. Specifically, since we show in (A.10) that *Y* = 1 at *τ* = 0 (equivalently *T* = *T*_*c*_) in the quasi-steady case, we introduce the delay times *τ*\* and *T*\*, defined through *Y* (*τ*\*) = 1 and 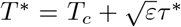, respectively. i.e. *T* = *T*\* is the time at which *Y* = 1. We refer to *T*\* as the dynamical position of the bifurcation. As we would expect, we have *T*\* = *T*_*c*_ for the quasi-steady case, which we identify above as *ω* ≪ *ε* ≪ 1 (from *λ* ≪ 1).

However, if *λ* ≫ 1 (equivalently *ε* ≪ *ω*), the intermediate dynamics are emphatically *not* quasi-steady, and we show below that 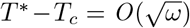. Importantly, this delay time is faster than the slow timescale (*T* = *O* (1)) and slower than the fast timescale (*T* = *O* (*ω*)), and arises because the bifurcation is crossed dynamically, even though this crossing timescale is slower than the fast reaction timescale.

In this case, in order for the large reaction rate term *λ*d*Y/*d*τ* to balance the self-repression/self-activation term *τY*, the appropriate intermediate timescale is actually 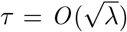, asymptotically larger than assumed for the generic intermediate timescale. We therefore introduce the rescaled intermediate timescale 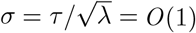, which is equivalent to 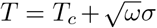 and 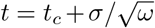. We emphasize that this suggests an appropriate intermediate timescale of 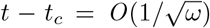 (equivalently 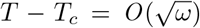) as the bifurcation is crossed, though we are able to be more precise below. Using this intermediate timescale 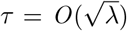, we also require that 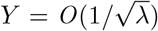 for consistency with the matching condition (A.9b). Hence, we also introduce *Y*_*L*_ = *O* (1) such that 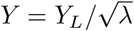. Using these *τ* and *Y* scalings in the system (A.9), we obtain the rescaled leading-order system

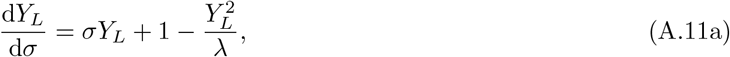

with appropriate matching condition

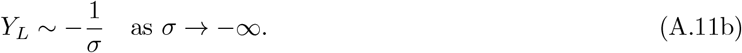

The leading-order version of (A.11) is

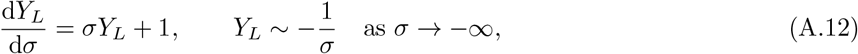

and is solved by

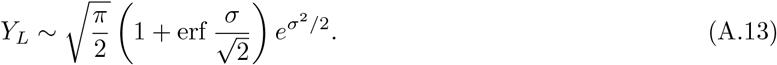

The solution (A.13) is only valid as long as the term neglected in going from (A.11) to (A.12) remains small. Therefore, the solution (A.13) is only valid for *σ < σ*_*c*_, where *σ*_*c*_ is to be determined. The quantity *σ*_*c*_ can be found formally by balancing the neglected 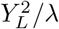 (saturation) term in (A.11) with the growing *σY*_*L*_ term. Noting that 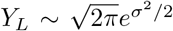 for large *σ*, balancing these terms asymptotically yields 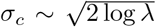. At this value of *σ*, we find that *Y*_*L*_ = *O*(*λ*), and hence that 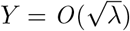, so the next appropriate balance is obtained by introducing *Y*_*R*_ such that 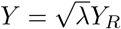. While it is relatively straightforward to solve the resultant leading-order system for *σ > σ*_*c*_ using these scalings (see e.g. Appendix B.1.2), the important result we require to calculate the dynamical position of the bifurcation is the value of *σ*\* such that *Y* (*σ*\*) = 1 (equivalently 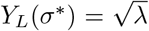). This will tell us the value of *T*\* via 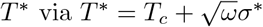. Given that (A.13) is valid until *Y*_*L*_ = *O* (*λ*), we may use (A.13) to deduce that 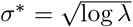, and hence that 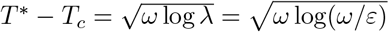.

To summarize, we have illustrated how the presence of a bifurcation in the toy system (A.6a) can lead to delayed dynamics over timescales of 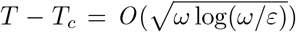 when *ε* ≪ *ω* ≪ 1. Importantly, these delay timescales are generally different to both fast (*T* = *O* (*ω*)) and slow (*T* = *O* (1)) timescales in the problem. This dynamic effect is sometimes referred to as ‘critical slowing down’, and is a universal feature of dynamical systems near bifurcations [30, 48, 57].

## B Asymptotic analysis of imperfect transcritical bifurcations

### B.1 Travelling-wave-like motion for off-to-on transcritical bifurcations

In this Appendix we consider imperfect transcritical pitchfork bifurcations, where the bifurcation moves unidirectionally from the patterned to the unpatterned state (i.e. off-to-on). The transcritical nature corresponds to the governing equation (6) with *n* = 2, and far-field conditions (7). The ‘off-to-on’ nature corresponds to *χ <* 0. To fully characterize the possible behaviours of the transition across the bifurcation, we seek to understand the possible behaviours of *Y* in terms of Λ and *χ*. We do this by examining the distinguished asymptotic limits of the ODE (6) with boundary conditions (7). This will allow us to derive analytic leading-order solutions, and subsequently to quantify the effect of dynamically crossing the bifurcation.

While the analytic results we obtain for *Y* and *Z*\* are specific to the case *χ <* 0, we note that in the case where *χ >* 0 (i.e. on-to-off), the asymptotic order of scalings for *Z*\* will be of the same order that we derive here for *χ <* 0. We proceed by analysing each region in increasing order of complexity.

#### B.1.1 Positive feedback dominates: Region A

The simplest region is Region A, where Λ, | *χ* | ≪ 1 and hence positive feedback dominates. In this case, the ODE (6) becomes quasi-steady:

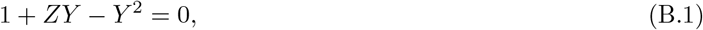

and is solved by

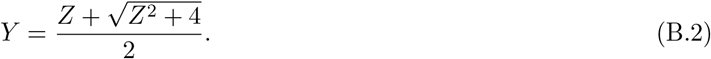

In this quasi-steady limit, *Y* (0) = 1, and therefore *Z*\* = 0. Hence, in Region A the bifurcation takes its equilibrium position as we would expect from a quasi-steady analysis, and the patterning is spatially local and quasi-steady.

#### B.1.2 Spatio-temporal variations dominate: Region B

Region B occurs when | *χ* | ≫ 1 and Λ*/* | *χ* | ^3*/*2^ ≪ 1. In this region, spatio-temporal changes in the parameters dominate and delaying effects become important, with implications for pattern robustness. Importantly, in this region there are sharp transitions between the unpatterned and patterned parts of the transition region, separated by *Z* = *Z*_*c*_ (which we shall see is different but related to *Z*\*).

To summarise the asymptotic structure of the solution before going into the specific mathematical analysis - in Region B, the interesting behaviour occurs over the lengthscale 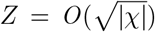, so we scale into a new independent variable 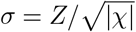. The asymptotic solution is then split into two asymptotic regions separated by the point 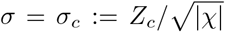. The determination of *σ*_*c*_ is a key goal of our analysis. The unpatterned region corresponds to *σ < σ*_*c*_ and the patterned region corresponds to *σ > σ*_*c*_. The solution is smaller in the unpatterned region, with the scaling 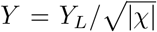, and larger in the patterned region, with the scaling 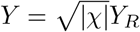.

Here, information travels in the same direction as the pattern transition i.e. from the unpatterned state to the patterned state (off-to-on). As such, our analysis starts in the unpatterned state, where *σ < σ*_*c*_. Using the scalings noted above, the leading-order scaled version of the ODE system (6), (7) in the unpatterned state (*σ < σ*_*c*_) is

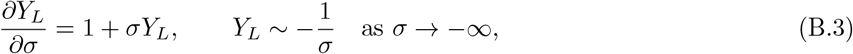

which is solved by

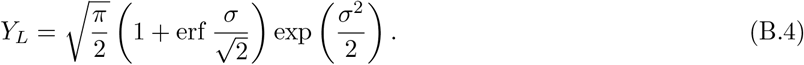

Given the relative scalings of *Y* in the unpatterned and patterned regions, we can determine *σ*_*c*_ by calculating when *Y*_*L*_ = *O* (|*χ*|). Since (B.4) yields the far-field result

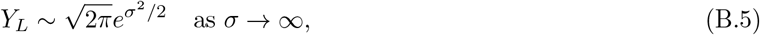

it is straightforward to note that 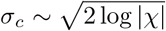.

For completeness, we also provide the solution in the patterned state, where *σ > σ*_*c*_. Using the scalings noted above for the patterned state, the leading-order scaled version of the ODE system (6), (7) in the patterned state (*σ > σ*_*c*_) is

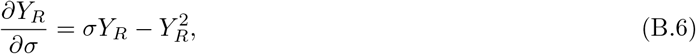

together with an appropriate matching condition with (B.4). The patterned system (B.6) is solved by

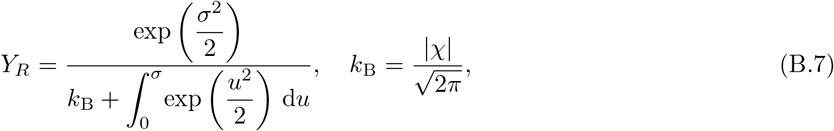

where *k*_B_ is a constant determined by matching with (B.4). The matching procedure involves the straightforward tracking of the exponentially growing term (B.5) within a logarithmic transition region where *Y*_*L*_ = *O* (|*χ*|). Specifically, an intermediate transition region where *σ* − *σ*_*c*_ = *O* (1*/σ*_*c*_).

Finally, we calculate the dynamic position of the bifurcation *<u>Z</u>* = *Z*\* at which *Y* (*Z*\*) = 1. Given the relative scalings of *Y* in each asymptotic region, we may infer 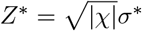 by calculating *σ*\* through the relationship 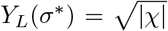. From (B.4), this occurs when 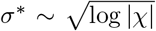, and hence we obtain the following asymptotic result for the dynamic position of the bifurcation:

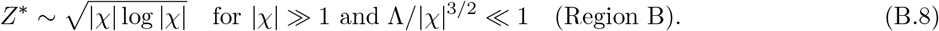

#### B.1.3 Diffusion dominates: Region C

Region C occurs when Λ ≫ 1 and Λ*/*| *χ*| ^3*/*2^ ≫1. In this region, diffusive effects dominate. In a similar manner as for Region B, there are sharp transitions between the unpatterned and patterned parts of the transition region here, separated by *Z* = *Z*_*c*_ (which again is different but related to *Z*\*).

To summarise the asymptotic structure of the solution - in Region C the interesting behaviour occurs over the lengthscale *Z* = *O* (Λ^1*/*3^), so we scale into a new independent variable *ζ* = *Z/*Λ^1*/*3^. The solution is again split into two asymptotic regions, matched using an intermediate transition region. This time, the two main asymptotic regions are separated by the point *ζ* = *ζ*_*c*_ := *Z*_*c*_*/*Λ^1*/*3^. The determination of *ζ*_*c*_ is a key goal of our analysis. The unpatterned region corresponds to *ζ < ζ*_*c*_ and the patterned region corresponds to *ζ > ζ*_*c*_. The solution is smaller in the unpatterned region, with the scaling *Y* = *W*_*L*_*/*Λ^1*/*3^, and larger in the patterned region, with the scaling *Y* = Λ^1*/*3^*W*_*R*_.

Using the scalings noted above, the leading-order scaled version of the system (6), (7) in the unpatterned state (*ζ < ζ*_*c*_) is

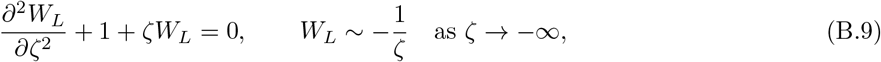

which is equivalent to the system (25), and hence solved by (26) for a different value of *c*_*L*_, to be determined through matching with the patterned region.

In the patterned state, where *ζ > ζ*_*c*_, the leading-order scaled version of the ODE (6), (7) is

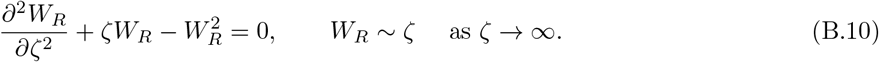

Given the relative scalings of *Y* in each asymptotic region, the position *ζ* = *ζ*_*c*_ can be obtained by calculating when *W*_*R*_ = *O* (Λ^−2*/*3^). Similarly, the dynamic position of the bifurcation *Z*\* = Λ^1*/*3^*ζ*\* can be inferred through the relationship *Y* (*Z*\*) = 1, and hence *W*_*R*_(*ζ*\*) = Λ^−1*/*3^. Hence, we require knowledge of the decaying behaviour of the nonlinear ODE (B.10) as we move towards the unpatterned far field *ζ* → −∞. Linearising (B.10) around the far-field trivial solution and imposing decay results in the behaviour

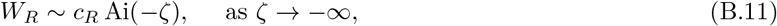

where *c*_*R*_ = *O* (1) is a constant that can be determined if required by solving (B.10) numerically and imposing decaying behaviour as *ζ* → −∞. Using the far-field behaviour of the Airy function, we find that *W*_*R*_ = *O* (Λ^−2*/*3^) for *ζ*_*c*_ ∼ − (log Λ)^2*/*3^ and that *W*_*R*_(*ζ*\*) = Λ^−1*/*3^ yields 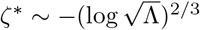, resulting in the following asymptotic result for the dynamic position of the bifurcation

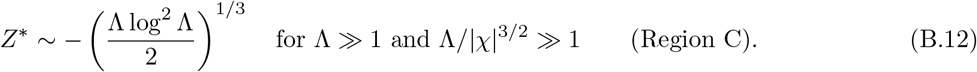

#### B.3 Spatio-temporal oscillations for transcritical bifurcations

Here, we consider spatio-temporal oscillations for transcritical bifurcations going from unpatterned-patterned-unpatterned. This is the transcritical equivalent of Section 3.2 in the main text. Essentially, we consider (1) with *n* = 2 and *a >* 0, which produces an imperfect transcritical bifurcation. As such, we start with the system

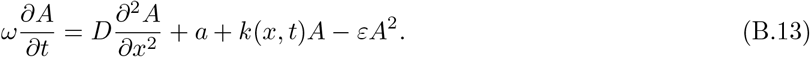

We are specifically interested in spatio-temporal oscillations of *k*(*x, t*) (i.e. cases where the bifurcation can turn around). As such, we follow the main text analysis here, and zoom into a region near the generic fixed point *x* = *x*\*, around the time *t* = *t*\* at which the bifurcation reaches the fixed point, defined through *k*(*x*\*, *t*\*) = 0. We consider the scenario where the point *x* = *x*\* transitions from unpatterned-patterned-unpatterned under a quasi-steady analysis (i.e. a local balance between reaction terms). We are specifically interested in understanding when the system overcomes the forcing to pattern and remains unpatterned in the dynamic case, demonstrating intrinsic robustness. Therefore, we investigate when the critical slowing down effect of Region B can overcome this forcing before the system returns to the locally patterned state. That is, quantifying when the system is robust to temporary incursions into regimes that would appear to be unpatterned from a quasi-steady analysis.

Since we are focusing on the critical slowing down scenario, diffusive effects can be neglected. As such, we can effectively consider *D* = 0 in (B.13) and in what follows. However, we retain *D* for the time being in order to understand when we can formally neglect it. As before, we treat *ω* and *ε* as small.

We are specifically interested in understanding when forcing transitions from unpatterned-patterned-unpatterned resist patterning and remain unpatterned, so the key terms in (B.13) are the left-hand side and the first two terms on the right-hand side. While we will formalize this intuitive dominant balance for the specific scenarios in which we are interested below, it is helpful to note that for this dominant balance all relevant asymptotic scalings will reduce (B.13) to an equation of the form

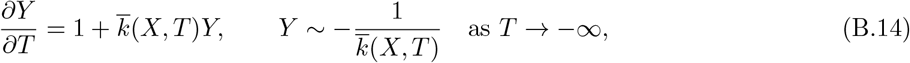

at leading-order, for appropriately defined independent variables *X, T*, dependent variable *Y* (*X, T*), and function 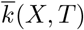, noting that 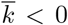 as *T* → −∞. Using the integrating factor method, it is straightforward to derive the following solution of the linear ODE (B.14) in terms of 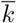:

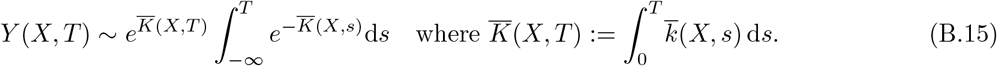

#### B.3 Fixed away from the turning point

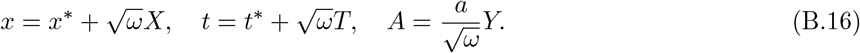

These scalings turn (B.13) into

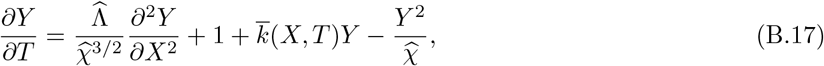

where we introduce the parameter groupings

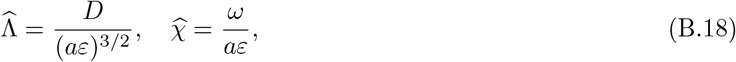

keeping similar notation as in (6b), and the function

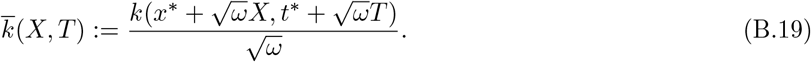

We note that 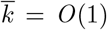 since *k*(*x*\*, *t*\*) = 0. Moreover, while it is possible to replace the right-hand side of (B.19) with its linearization for many differentiable functions *k*, we keep it in its general form since we are specifically interested in oscillations in *T*. That is, we are specifically interested in 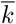 with turning points in *T*, and these will generally not be well-approximated by their linearisations.

The formal neglect of the diffusive and saturation terms in (B.17) correspond to the limits 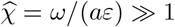 and 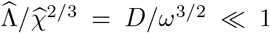, directly analogous to the limits that take us into Region B in the previous subsection. Moreover, these limits reduce (B.17) to the system (B.14), which is solved by (B.15).

For any specified oscillatory function *k*(*x, t*) (and hence 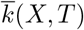), we can calculate when *Y* reaches a critical minimal value (specified by the particular problem being considered) via 1D root finding of the analytic solution (B.15).

### B.4 Fixed at the turning point

We now consider the case where *k*_*t*_(*x*\*, *t*\*) = 0 and *k*_*tt*_(*x*\*, *t*\*) *<* 0 i.e. the bifurcation turns around at the fixed point *x* = *x*\*. In this case, the appropriate inner equation scalings into the initially unpatterned region are

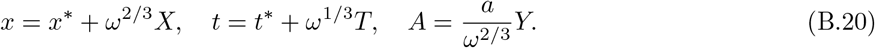

These scalings turn (B.13) into

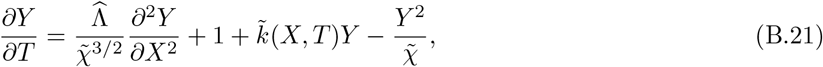

where 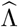 is defined in (B.18), and we introduce the parameter grouping

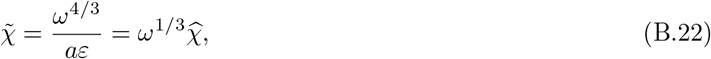

where 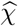 is defined in (B.18), as well as the function

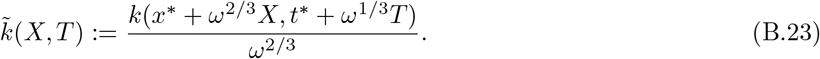

The formal neglect of the diffusive and saturation terms in (B.21) correspond to the limits 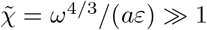 and 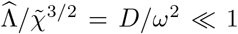. If these constraints are satisfied, this limit reduces (B.21) to the system (B.14), which is solved by (B.15).

## C Biological example: bacterial quorum sensing

### C.1 Model

We consider bacterial quorum sensing (QS) in *Vibrio fischeri*, which causes bioluminescence in the Hawaiian bobtail squid [36]. We model the LuxR quorum sensing system under the influence of two autoinducers, 3OC6HSL and C8HSL, with cross-talk between them [58]. We model the concentration *Ã* of one of the autoinducers, 3OC6HSL, which is part of a positive feedback loop: it forms a complex with a cognate protein LuxR that activates the expression of LuxI, a synthase that catalyses the expression of 3OC6HSL. We refer to 3OC6HSL as AI here for brevity.

The dynamic dimensional governing equations inside the two-dimensional cell layer with length 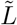 and height 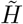 are

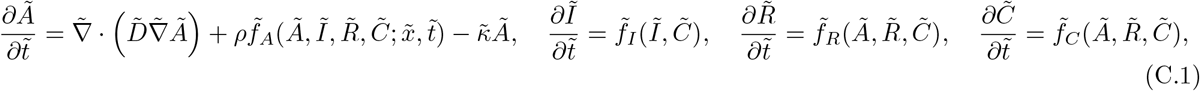

For 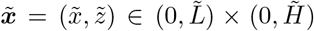, where 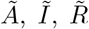, and 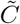 are the concentrations of autoinducer, LuxI, LuxR, and autoinducer-LuxR complex, respectively, within the cell layer. Here, 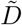 is the diffusivity of autoinducer within the cell layer, *ρ* is the volume fraction of cells-to-total-volume, 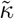 is the decay rate of autoinducer, and the reaction terms 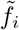 for *i* ∈ *{A, I, R, C}* (which occur within each bacterium) are defined as the following:

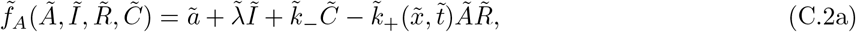

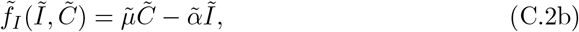

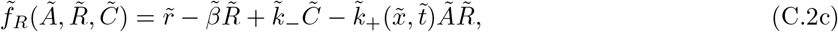

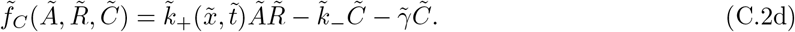

There is cross-talk with the second autoinducer, C8HSL, via competitive binding to the cognate protein LuxR. We account for this effect though a space- and time-dependent AI-LuxR binding parameter 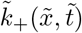. This models spatio-temporal oscillations in C8HSL, which could be caused by growth or motility of a competing bacterial population, changes in gene-expression, fluctuations in mass transport (which drive autoinducer concentration gradients [12]), or a spatio-temporally fluctuating signal in a synthetic system. We note that our model assumes that the C8HSL-LuxR complex does not promote the expression of LuxR, in line with experimental findings [58]. Specifically, we let

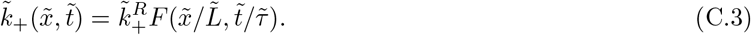

We will generally use either

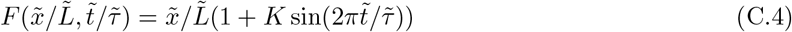

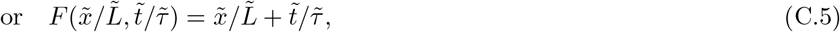

where we assume a linear spatial gradient in the binding strength for simplicity. We apply no flux boundary conditions on the sides and bottom of the domain,

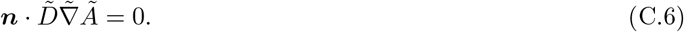

and a mixed boundary condition on the upper boundary,

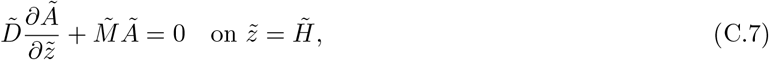

where 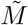 is a mass transfer coefficient that models mass transfer via a fluid flow or diffusion into an open space surrounding the population, as is common in natural environments and microfluidic experiments [12]. As initial condition, we use the steady solution of the system for 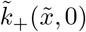.

Typical orders of magnitude of the dimensional parameters in this problem are estimated in Table 2.

### C.2 Dimensionless problem

We scale

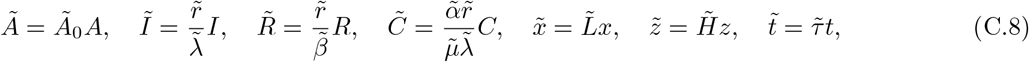

where the metabolite concentrations have been scaled with the typical values they take in the unpatterned region, where

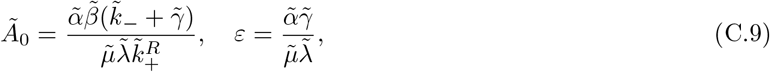

and where time has been rescaled with the dynamic forcing mechanism in our system: the timescale of kinetic variation.

Inside the cell layer, (C.1) becomes

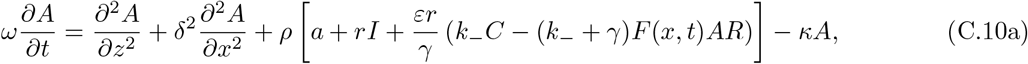

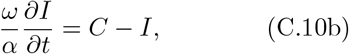

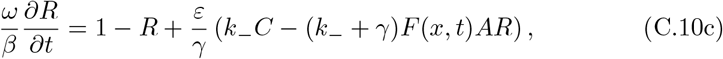

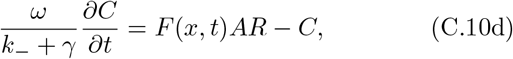

and we have introduced the dimensionless parameters

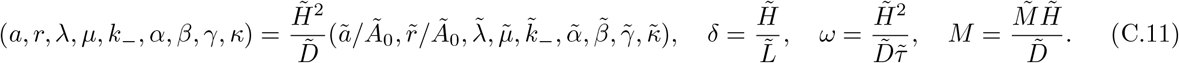

The nondimensional versions of the boundary conditions remain no flux on the sides and bottom of the domain,

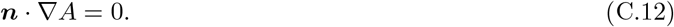

and a mixed boundary condition on the upper boundary,

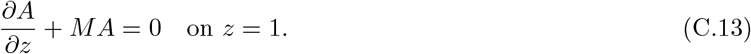

We proceed by exploiting the small parameters 0 *< ε, δ, ω*≪ 1, treating all other dimensionless parameters as *O* (1).

### C.3 Solution in unpatterned region

Taking the limits *ε, δ, ω* → 0, in the unpatterned regime (whose extent will become clear shortly) where *A, I, C, R* = *O* (1), the leading-order version of (C.10) is linear, and its solution is

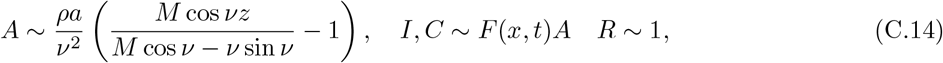

where *ν* = *ν*(*x, t*) is defined through

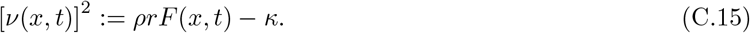

The extent of the unpatterned region can be determined through defining the moving boundary *s*(*t*), defined implicitly through

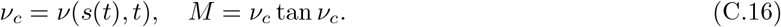

The unpatterned region is the set of (*x, t*) satisfying *ν*(*x, t*) *< ν*_*c*_.

### C.4 Derivation of inner problem

We define the inner transition region through the inner scalings

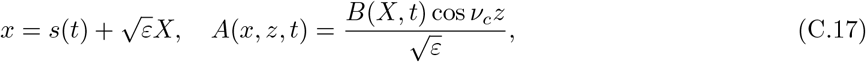

noting that 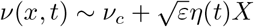 in the inner region, where

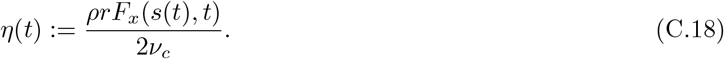

After some inner analysis involving the derivation of a solvability condition at next asymptotic order (similar to that in [12]), we obtain the following governing equation for the scaled amplitude *B*:

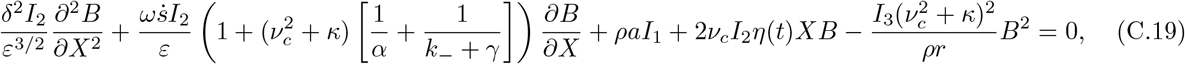

with boundary conditions

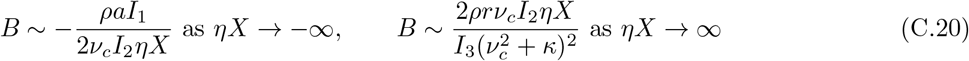

Where 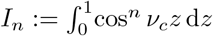, and hence

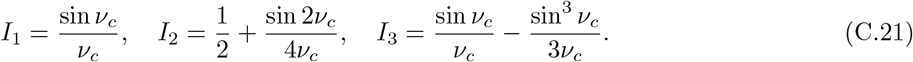

In deriving (C.19), we have implicitly considered the distinguished asymptotic limit where *δ* = *O* (*ε*^3^) and *ω* = *O* (*ε*). Subcases which do not satisfy these precise scalings can be distilled as sublimits of the distinguished limit we consider.

### C.5 Reduction to canonical form

The governing equation (C.19) takes the general form:

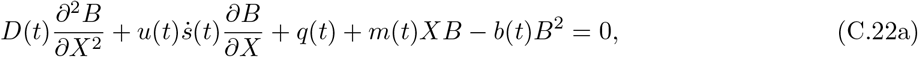

where *D, q, b >* 0, and with boundary conditions

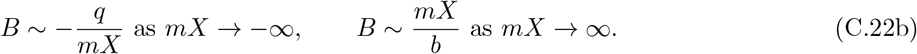

Broadly, (C.22) is the general form we would expect for an imperfect transcritical bifurcation. Using the transformation variables

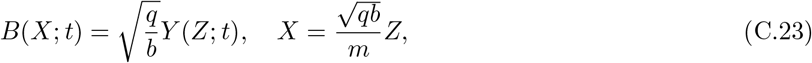

the equation (C.22) is transformed into

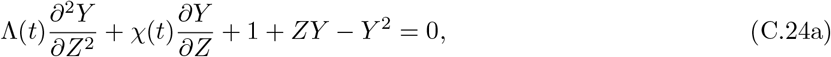

with boundary conditions

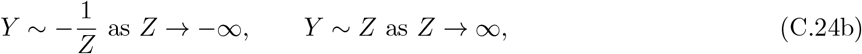

which is precisely the general form we derive in (6a), (7). This time, however, the coefficients Λ(*t*) and *χ*(*t*) are given as

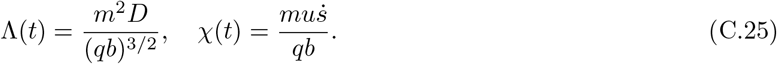

In terms of the quorum sensing system presented in (C.19), we have:

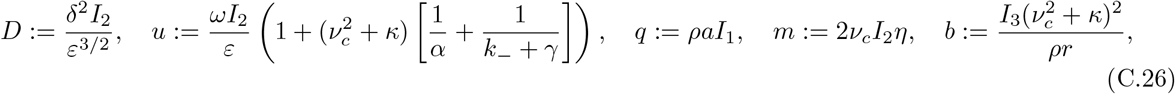

and so

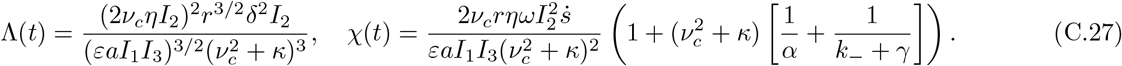

### C.6 Oscillating parameters

We now consider the case when the bifurcation motion can turn around. To analyse and quantify this effect, we scale into the region close to a generic fixed point *x* = *x*\*, around the time *t* = *t*\* at which the bifurcation reaches this point. That is, *t*\* is defined implicitly through *ν*(*x*\*, *t*\*) = *ν*_*c*_.

We are specifically interested in the critical slowing down region where delayed effects dominate. We therefore use the new length, time, and concentration scalings:

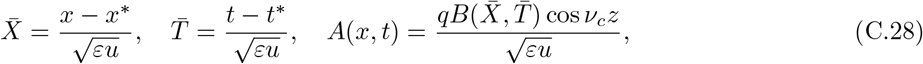

A similar inner analysis to before yields the general leading-order amplitude equation

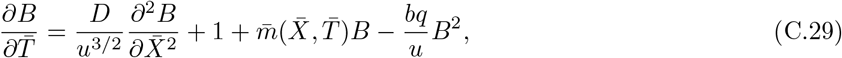

where the parameters take the same values as in (C.26), except now with

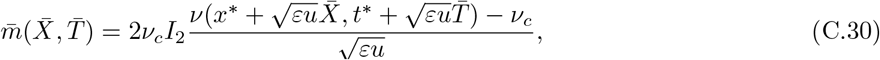

where *ν*_*c*_ is defined in (C.16). Then we see that if *u*^3*/*2^ ≫ *D* (formally *ω* ≫ *δ*^4*/*3^) and *u* ≫ 1 (formally *ω* ≫ *ε*), the system (C.29) reduces to an equation of the form (B.14), which is solved by (B.15).

Hence, for a specified function *F* (*x, t*) (and hence *ν*(*x, t*), and hence 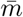 in (C.30)), we can calculate the value of *B* through the general solution (B.15). Moreover, for finite domains *x* ∈ (0, 1), if we apply this analysis at the boundary *x* = 1,^3^ we can calculate the critical oscillation period for which patterns are suppressed (i.e. remain below a specified critical value) via simple 1D root finding (see Fig. 4).

## Footnotes

1 For example, for a Turing bifurcation activated through a pitchfork with *n* = 3, *A* typically represents the envelope amplitude for the patterned state, as predicted by the fastest growing mode at the onset of instability.

2 Assuming that our analysis holds near a boundary, which should hold unless the amplitude varies rapidly i.e. unless the system exhibits a boundary layer near *x* = 1.

3 Which should hold as long as there is no boundary layer near *x* = 1.

## Notes

### Competing Interest Statement

The authors have declared no competing interest.

### Summary of Updates

Supplemental information incorporated into main text and appendices

